# Synaptic and intrinsic mechanisms impair reticular thalamus and thalamocortical neuron function in a Dravet syndrome mouse model

**DOI:** 10.1101/2021.09.03.458635

**Authors:** Carleigh Studtmann, Marek Ladislav, Mackenzie A. Topolski, Mona Safari, Sharon A. Swanger

**Author notes:** **Corresponding Author:** Sharon A. Swanger, Fralin Biomedical Research Institute at VTC, Virginia Tech, 2 Riverside Circle, Roanoke, VA 24016.

## Abstract

Thalamocortical network dysfunction contributes to seizures and sleep deficits in Dravet syndrome (DS), an infantile epileptic encephalopathy, but the underlying molecular and cellular mechanisms remain elusive. DS is primarily caused by mutations in the *SCN1A* gene encoding the voltage-gated sodium channel Na_V_1.1, which is highly expressed in GABAergic reticular thalamus (nRT) neurons as well as glutamatergic thalamocortical neurons. We hypothesized that Na_V_1.1 haploinsufficiency alters somatosensory corticothalamic circuit function through both intrinsic and synaptic mechanisms in nRT and thalamocortical neurons. Using *Scn1a* heterozygous mice of both sexes aged P25-P30, we discovered reduced intrinsic excitability in nRT neurons and thalamocortical neurons in the ventral posterolateral (VPL) thalamus, while thalamocortical ventral posteromedial (VPM) neurons exhibited enhanced excitability. Na_V_1.1 haploinsufficiency enhanced GABAergic synaptic input and reduced ascending glutamatergic sensory input to VPL neurons, but not VPM neurons. In addition, glutamatergic cortical input to nRT neurons was reduced in *Scn1a* heterozygous mice, whereas cortical input to VPL and VPM neurons remained unchanged. These findings introduce input-specific alterations in glutamatergic synapse function and aberrant glutamatergic neuron excitability in the thalamus as disease mechanisms in Dravet syndrome, which has been widely considered a disease of GABAergic neurons. This work reveals additional complexity that expands current models of thalamic dysfunction in Dravet syndrome and identifies new components of corticothalamic circuitry as potential therapeutic targets.

**HIGHLIGHTS:**

- GABAergic reticular thalamus neurons have impaired tonic and burst firing properties in a Na_V_1.1 haploinsufficiency mouse model of Dravet syndrome.
- Na_V_1.1 haploinsufficiency has opposing effects on spike firing in two distinct glutamatergic thalamocortical neuron populations.
- Na_V_1.1 haploinsufficiency alters glutamatergic synaptic connectivity in an input-specific manner in the thalamus.
- Dysregulation of both intrinsic and synaptic mechanisms contribute to imbalanced thalamic excitation and inhibition in this Dravet syndrome mouse model.

## INTRODUCTION

Dravet syndrome (DS) is an infantile epileptic encephalopathy most often caused by mutations in the *SCN1A* gene, which encodes the alpha subunit of the Na_V_1.1 voltage-gated sodium channel (Claes et al., 2001; Dravet, 2011). DS is characterized by intractable convulsive and non-convulsive (absence) seizures in infancy as well as progressive symptomology including ataxia, intellectual disability, attention deficits, autistic features, sleep disruptions, and a high risk of sudden death (Berkvens et al., 2015; Darra et al., 2019; Dravet, 2011; Licheni et al., 2018; Ragona, 2011; Rodda et al., 2012; Takayama et al., 2014). Na_V_1.1 is expressed in the axon initial segment as well as distal axons of primarily, but not exclusively, GABAergic neuron populations (Favero et al., 2018; Hedrich et al., 2014; Ogiwara et al., 2007). More than half of DS mutations lead to haploinsufficiency of Na_V_1.1, which disrupts the excitatory/inhibitory balance across a variety of brain circuits in DS models (Bender et al., 2016; Kalume et al., 2007; Ogiwara et al., 2007; Ritter-Makinson et al., 2019; Tai et al., 2014). The multifaceted DS phenotype likely arises from dysfunction of specific cell-types within these brain circuits (Bender et al., 2013; Han et al., 2012; Hedrich et al., 2014; Kalume et al., 2015; Kalume et al., 2007; Ogiwara et al., 2007; Ritter-Makinson et al., 2019; Rubinstein et al., 2015a; Tai et al., 2014), which may contribute to the limited efficacy of treatments targeting brain-wide excitation or inhibition in DS (Chiron, 2011; Cross et al., 2019; Gataullina and Dulac, 2017; Takayama et al., 2014). Therefore, elucidating the precise molecular and cellular mechanisms underlying dysfunction in particular circuits may provide more specific therapeutic targets to ameliorate DS symptoms.

The somatosensory corticothalamic (CT) circuit controls somatic information flow between the periphery and the cerebral cortex, and it is involved in attention, pain processing, and sleep (Beenhakker and Huguenard, 2009; Wolff and Vann, 2019; Zikopoulos and Barbas, 2006). This circuit generates oscillatory sleep spindles that are critical for NREM sleep (Fernandez and Luthi, 2020), and under pathological conditions these oscillations can become hypersynchronous, leading to absence seizures (Cheong and Shin, 2013; Lee et al., 2004; McCafferty et al., 2018). Altered thalamic activity is evident in some DS patients (Moehring et al., 2013), and DS mouse models exhibit somatosensory CT circuit dysfunction including reduced sleep spindles, hypersynchronous oscillations, and absence seizures (Hedrich et al., 2014; Kalume et al., 2015; Ritter-Makinson et al., 2019). However, the molecular and cellular mechanisms underlying thalamic dysfunction in DS are not well understood.

The somatosensory CT circuit comprises the somatosensory cortex, the ventrobasal (VB) thalamus, and the reticular nucleus of the thalamus (nRT). The VB thalamus includes two glutamatergic thalamocortical neuron populations - the ventral posterolateral (VPL) and ventral posteromedial (VPM) thalamus. VPL and VPM neurons receive distinct ascending somatosensory information and transmit it to the somatosensory cortex, which provides reciprocal glutamatergic feedback to the thalamus (Ab Aziz and Ahmad, 2006; Brecht and Sakmann, 2002; Lenz, 1992). These corticothalamic and thalamocortical projections send axon collaterals to the GABAergic nRT, which provides the primary inhibitory input to the VPL and VPM (**Figure 1A**). Neurons in the nRT, VPL, and VPM have two firing modes critical for somatosensory CT circuit function: a tonic firing mode involved in somatosensory processing, and a burst firing mode underlying intra-thalamic oscillations (Sherman, 2001; Steriade and Llinas, 1988). GABAergic nRT neurons as well as glutamatergic VPL and VPM neurons express Na_V_1.1 at high levels (Ogiwara et al., 2013; Papale et al., 2013). Interestingly, many glutamatergic neuron populations do not highly express Na_V_1.1, so its expression in VPL and VPM neurons may uniquely contribute to disease pathophysiology and provide distinctive therapeutic targets (Ogiwara et al., 2013).

**Figure 1.**
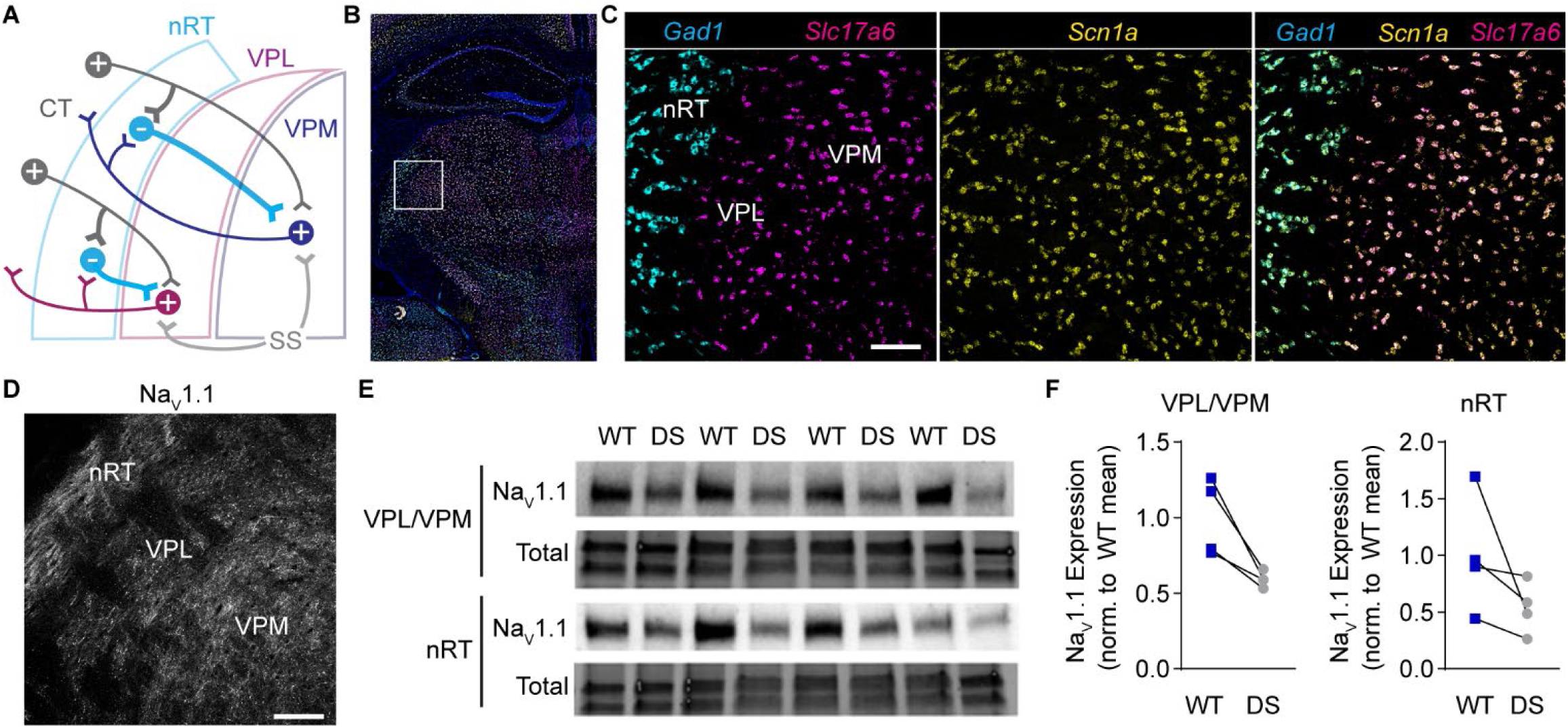
*Scn1a* mRNA and NaV1.1 protein expression in the somatosensory thalamus. **A**. A circuit diagram illustrates somatosensory corticothalamic (CT) circuit connectivity. Glutamatergic CT neurons innervate nRT, VPL, and VPM neurons. GABAergic nRT neurons innervate VPL and VPM neurons, which send glutamatergic projections to the cortex and collaterals to the nRT. Distinct ascending glutamatergic somatosensory (SS) afferents innervate VPL and VPM neurons. **B**. A representative 10X tiled image of a coronal mouse brain section shows *Scn1a* (yellow), *Gad1* (cyan), and *Slc17a6* (magenta) mRNA labeled by FISH with DAPI counterstain (blue). **C**. 20X images of the boxed region in panel B show *Gad1*+ and *Slc17a6*+ neurons express *Scn1a* mRNA. **D**. A representative 20X image shows NaV1.1 immunolabeling in the nRT, VPL, and VPM. Scale bar: 100 μm (**C,D**). **E**. A western blot shows NaV1.1 and total protein expression in the nRT and VPL/VPM for WT and DS mice (n = 4 littermate pairs). **F**. NaV1.1 protein levels were quantified by densitometry and normalized to the WT mean. Individual data points represent each mouse.

Given the delicate balance of excitation and inhibition in the somatosensory CT circuit, changes in either intrinsic excitability or synaptic connectivity could significantly alter thalamocortical network function in DS. To effectively target thalamic dysfunction, it is critical to understand precisely how the circuitry is dysregulated. We hypothesized that Na_V_1.1 haploinsufficiency alters somatosensory CT circuit function through both intrinsic and synaptic mechanisms in GABAergic nRT and glutamatergic thalamocortical neurons. Herein, we report cell-type- and synapse-specific alterations in nRT and thalamocortical neurons that expand current models of somatosensory thalamus dysfunction in DS by introducing glutamatergic neuron excitability and thalamic synapse dysfunction as novel disease mechanisms.

## MATERIALS AND METHODS

### Mouse model

Mouse studies were performed according to protocols approved by the Institutional Animal Care and Use Committee at Virginia Polytechnic Institute and State University and in accordance with the National Institutes of Health guidelines. *Scn1a*^*tm1Kea*^ mice (037107-JAX), which are on a 129S6/SvEvTac background, were purchased from the Mutant Mouse Resource and Research Center (Miller et al., 2014). The *Scn1a*^*tm1Kea*^ colony was maintained by crossing heterozygous 129S6/SvEvTac-*Scn1a*^*tm1Kea/WT*^ mice with wild type 129S6/SvEvTac (WT) mice. Experimental mice were F1 hybrids generated by crossing heterozygous 129S6/SvEvTac-*Scn1a*^*tm1Kea/WT*^ mice with C57BL/6J mice (000664, JAX). Genotyping was performed by Transnetyx using real-time PCR. Mice were housed in a 12 hour light/dark cycle with *ad libitum* access to food and water. Age-matched WT and *Scn1a*^+/-^ littermates of both sexes aged P25-P30 were used for all experiments.

### Fluorescence in situ hybridization

For mRNA detection, mice were euthanized by isoflurane overdose and brains were rapidly dissected and flash frozen in OCT. Brains were cryosectioned (20 μm), immediately fixed with 4% paraformaldehyde, and RNAscope (ACDBio) multiplex fluorescent in situ hybridization (FISH) was performed according to the manufacturer’s instructions with probes against *Scn1a, Gad1, and Slc17a6* (mRNA for VGLUT2). A tiled 10X image was acquired with CellSens software and an Olympus IX83 widefield fluorescence microscope equipped with a Hamamatsu Orca Flash IV camera, X-Cite Xylis LED, and DAPI, FITC, TRITC, and Cy5 filter sets. 20X images were acquired with a Zeiss 700 confocal and Zen Black software using 405, 488, 533, and 633 lasers. The images were processed for presentation using ImageJ software.

### Western blotting

Mice were euthanized by isoflurane overdose and transcardially perfused with ice-cold 1X PBS. Brains were dissected and coronal slices (1 mm) were made using a brain matrix. Tissue punches from the VB thalamus and nRT were snap frozen in liquid nitrogen. Tissue was lysed with a sonicator in buffer containing (in mM) 65 Tris base, 150 NaCl, 1 EDTA, 50 NaH_2_PO_4,_ 10 Na_4_P_2_O_7_, 1 Na_3_VO_4,_ 50 NaF, 1% Triton X-100, and 0.5% deoxycholic salt. Samples were centrifuged at 13,500 x *g* for 15 min at 4°C, denatured using Laemmli sample buffer with 200 mM dithiothreitol, and heated at 95°C. Equal amounts of protein were run on Mini-PROTEAN TGX stain-free gels, which were activated using UV light exposure to covalently bind fluorophores to protein. Protein was transferred to polyvinylidene difluoride (PVDF) membranes, and total protein content was imaged on a ChemiDoc MP System. Membranes were probed with anti-Na_V_1.1 antibodies (1:1000; UC Davis/NIH NeuroMab Facility; K74/71) and goat anti-mouse HRP-conjugated secondary antibodies (1:10,000; Jackson Immunoresearch; 115-035-146), and then visualized with Pico Plus substrate (ThermoFisher) and a ChemiDoc MP System. Bands were quantified using Image Lab, and band intensity was normalized to total protein levels.

### Immunohistochemistry

Na_V_1.1 immunohistochemistry was performed as previously described (Alshammari et al., 2016). Briefly, mice were euthanized by isoflurane overdose and transcardially perfused with 1X PBS followed by ice-cold 1% paraformaldehyde (PFA) and 0.5% methanol in 1X PBS. Brains were post-fixed in 1% PFA and 0.5% methanol in 1X PBS for 24 hr, transferred to 30% sucrose in 1X PBS for 48 hr, and then flash frozen in OCT. Cryosections (20 μm) were slide-mounted, treated with 100% acetone at −20°C for 7 min, blocked with FAB fragments and 10% normal donkey serum (NDS), and then immunostained with mouse anti-Na_V_1.1 (1:200; UC Davis/NIH NeuroMab Facility; K74/71) and Alexa 488-conjugated anti-mouse IgG1 secondary antibodies (ThermoFisher). Images were acquired with a Zeiss 700 confocal, 20X objective, 488 argon laser, and Zen Black software.

For synaptic marker immunostainings, mice were transcardially perfused with 1X PBS followed by 4% PFA in 1X PBS, and the brains were dissected, post-fixed in 4% PFA for 24 hr, cryoprotected in 30% sucrose in 1X PBS, and flash frozen in OCT. Cryosections (20 μm) were treated with 0.8% sodium borohydride in 1X Tris-buffered saline (TBS) at room temperature, and then 0.01M sodium citrate buffer, pH 6.0, at 100°C for 10 min. Sections were permeabilized with 0.5% Triton X-100 in 1X TBS, pH 7.4, and immunostained with antibodies against: VGLUT1 (1:100; Synaptic Systems; 135-511), VGLUT2 (1:200; Synaptic Systems; 135-402), VGAT (1:200; Synaptic Systems; 131-103), and gephyrin (1:200; Synaptic Systems; 147-011). Sections were incubated with Alexa 555-conjugated anti-mouse IgG1 and Alexa 488-conjugated anti-rabbit secondary antibodies (ThermoFisher), as appropriate. Images were acquired with a Zeiss 700 laser-scanning confocal, a 100X objective, and 488 or 533 lasers. Three random fields were imaged from the left and right nRT, VPL, and VPM in two sections for a total of 9 – 12 images per mouse.

### Image Analysis

Images were assigned numerical identifiers and analysis was performed blinded to genotype. The *Synapse Counter* plugin from ImageJ was used to analyze the number and size of VGAT/gephyrin, VGLUT1, and VGLUT2 puncta. Synapse size restrictions in *Synapse Counter* were set as follows: VGAT in all regions (20-1000 px^2^), gephyrin in all regions (10-400 px^2^), VGLUT1 in all regions (10-400 px^2^), VGLUT2 in nRT (70-1200 px^2^), VGLUT2 in VPL and VPM (60-3000 px^2^). Data from each synapse marker were averaged for all images from one mouse. The area occupied by cell bodies and white matter tracks were automatically detected in VGLUT1 images using ImageJ, and the number of synapses was corrected for this area. Images were processed for presentation in ImageJ, and WT and DS images within each figure are displayed with equivalent intensity settings.

### Electrophysiology

Mice were deeply anesthetized with an overdose of inhaled isoflurane and transcardially perfused with ice-cold sucrose-based artificial cerebrospinal fluid (aCSF) containing (in mM) 230 sucrose, 24 NaHCO_3_, 10 glucose, 3 KCl, 10 MgSO4, 1.25 NaH_2_PO_4_, and 0.5 CaCl_2_ saturated with 95% O_2_ / 5% CO_2_. The brain was removed and glued to a vibratome stage (Leica VT1200S), and horizontal 300 μm slices were cut in an ice-cold sucrose-aCSF bath. Slices were incubated in a NaCl-based aCSF containing (in mM) 130 NaCl, 24 NaHCO_3_, 10 glucose, 3 KCl, 4 MgSO4, 1.25 NaH_2_PO_4_, and 1 CaCl_2_ saturated with 95% O_2_ / 5% CO_2_ at 32°C for 30 min. The slices were then equilibrated to room temperature (RT) for 30 min and maintained at RT until used for recordings up to 8 hr later.

Recordings were made using a Multiclamp 700B amplifier (Molecular Devices), sampled at 20 kHz (Digidata 1550B, Molecular Devices), and low-pass filtered at 10 kHz using Axon pClamp 11 software (Molecular Devices). The extracellular recording solution contained (in mM) 130 NaCl, 24 NaHCO_3_, 10 glucose, 3 KCl, 1 MgSO4, 1.25 NaH_2_PO_4_, and 2 CaCl_2_ saturated with 95% O_2_ / 5% CO_2_, and was maintained at 32°C for recordings. For whole-cell current-clamp recordings, borosilicate glass recording electrodes (4-5 MΩ) were filled with (in mM) 130 K-gluconate, 4 KCl, 2 NaCl, 10 HEPES, 0.2 EGTA, 4 ATP-Mg, 0.3 GTP-Tris, 14 phosphocreatine-K, and 0.1% biocytin, pH 7.3. Pipette capacitance neutralization and bridge balance were enabled during current-clamp recordings for capacitance and series resistance compensation. To analyze intrinsic membrane properties, voltage responses were elicited by 200 ms hyperpolarizing current injections between 20 – 100 pA (20 pA steps). Depolarization-induced spike firing was elicited by 500 ms depolarizing current injections between 10 – 400 pA (10 pA steps). Hyperpolarization-induced rebound bursting was elicited by 500 ms hyperpolarizing current injections between 50 – 200 pA (50 pA steps). Three trials were completed for each current injection experiment.

For whole-cell voltage-clamp recordings, borosilicate glass recording electrodes (4-5) MΩ were filled with (in mM) 120 CsMeSO3, 15 CsCl, 8 NaCl, 10 tetraethylammonium chloride, 10 HEPES, 1 EGTA, 3 Mg-ATP, 0.3 Na-GTP, 1.5 MgCl2, 1 QX-314, and 0.1% biocytin, pH 7.3. After a ten minute equilibration period, mEPSCs and mIPSCs were recorded in the presence of 1 μM TTX in two minute epochs at a holding potential of −70 mV and 0 mV, adjusted for the liquid junction potential, respectively. Series resistance and cell capacitance were monitored throughout the experiment, but were not compensated, and cells were excluded if either parameter changed >20%.

### Biocytin labeling in acute brain slices

To confirm the cell location after electrophysiology recordings, brain slices were fixed with 4% paraformaldehyde in 1X PBS, pH 7.4, overnight at 4°C, washed in 1X PBS, and then stored at −20°C in cryoprotectant solution containing 0.87 M sucrose, 5.37 M ethylene glycol, and 10 g/L polyvinyl-pyrrolidone-40 in 0.1 M phosphate buffer, pH 7.4. For biocytin labeling, slices were blocked with 10% NDS in 1X PBS with 0.25% Triton X-100, and then incubated with 1.0 μg/ml DyLight 594-conjugated streptavidin (Jackson Immunoresearch) in blocking solution overnight at RT. Slices were stained with DAPI for 1 hr at RT and placed on a glass slide with a coverslip and DABCO mounting media (Sigma Aldrich), which was sealed with nail polish. 10X images were acquired on an Olympus IX83 microscope with a Hamamatsu Orca Flash 4.0 camera, X-Cite Xylis LED, and a TRITC filter set using CellSens software.

### Electrophysiology data analysis

Recordings were assigned numerical identifiers and analysis was performed blind to genotype. All current-clamp recordings were analyzed in Clampfit 11 (Molecular Devices). Resting membrane potential (RMP) was measured from current-clamp recordings immediately upon breakthrough of the cell membrane. Reported RMP values were corrected for the liquid junction potential. Input resistance was determined from the amplitude of voltage responses to 200 ms hyperpolarizing current injections, and the time constant and cell capacitance were determined by a mono-exponential fit of the voltage response. The Clampfit 11 threshold detection module was used to quantify the number of spikes and latency to the first spike in response to 500 ms depolarizing current injections or upon removal of 500 ms hyperpolarizing current injections. Rheobase was defined as the smallest depolarizing current injection that elicited an action potential. Values were averaged across three runs for all current injection experiments.

For mIPSC and mEPSC recordings, the six minutes following the equilibration period were analyzed to determine inter-event interval, amplitude, and decay kinetics using MiniAnalysis software (Synaptosoft). Data were digitally filtered at 1 kHz. Automated detection identified events with amplitude ≥ 7 pA (∼5 x RMS noise), and then events ≥ 5 pA were manually detected and automated detection accuracy was assessed. The 70-30% decay time was utilized to group mEPSC events as: Type 1 (decay time < 1 ms) or Type 2 (decay time > 1 ms), with this critical value determined by analyzing the decay time frequency histogram. Event data within each group were averaged for each cell and used for group data reporting and comparisons. Events within each group from all cells were combined for cumulative distribution plots. Reported decay times are weighted time constants determined from a two-component exponential fit of Type 1 mEPSC, Type 2 mEPSC, or mIPSC ensemble averages from each cell.

### Statistical analysis

*A priori* power analyses were performed in GPower 3.1 to estimate required samples sizes given appropriate statistical tests with α= 0.05, power (1 – β) = 0.8, and a moderate effect size or effect sizes based on pilot data. Statistical analyses were performed in GraphPad Prism version 9.0.2. All datasets were tested for normality with the Shapiro-Wilk test and equal variances using an F test (two independent groups) or the Brown-Forsythe test (three or more groups). Normal datasets with equal variances were analyzed by unpaired t-test or ANOVA with corrections for multiple comparisons. Non-normal datasets were analyzed with the Mann-Whitney U test (two independent groups). Repeated measures datasets were analyzed by two-way ANOVA, and a mixed-effects analysis was used when any data points were missing from repeated measures designs. The specific statistical test used as well as the test statistics are reported in the respective figure legend or table. All group data are plotted as mean ± s.e.m. in the figures, and numerical data reported in the text are mean ± s.e.m.

## RESULTS

### Scn1a mRNA and Na_V_1.1 protein expression in GABAergic and glutamatergic neurons in the thalamus

To evaluate how thalamic neuron function is impaired in DS, we utilized the F1 hybrid 129S-*Scn1a*^+/-^ x C57BL/6J mouse model at age P25-P30, hereafter referred to as DS mice. Given extensive strain differences in DS mouse model phenotypes, we first examined *Scn1a* mRNA and Na_V_1.1 protein levels in the thalamus to confirm high expression in both GABAergic and glutamatergic neurons in WT (F1 hybrid 129S-*Scn1a*^*+/+*^ x C57BL/6J) mice at P25-P30. Indeed, multiplex FISH revealed abundant *Scn1a* mRNA expression in *Gad1*-positive GABAergic nRT neurons as well as glutamatergic VPL and VPM neurons detected by *Slc17a6* mRNA, which encodes VGLUT2 (**Figure 1B,C**). Na_V_1.1 protein was detected in the nRT, VPL, and VPM by immunohistochemistry (**Figure 1D**). In addition, western blotting of VPL/VPM and nRT tissue punches showed reduced Na_V_1.1 protein levels in DS mice relative to WT littermates (VPL/VPM: 59.4 ± 2.6% of WT; nRT: 53.8 ± 1.1% of WT; **Figure 1E,F**), confirming haploinsufficiency in the thalamus of this DS model.

### Impaired GABAergic nRT neuron excitability in DS mice

Previous studies have reported opposing effects on nRT neuron excitability in different DS mouse models as well as age-dependent changes in GABAergic neuron excitability in the cortex (Favero et al., 2018; Hedrich et al., 2014; Kalume et al., 2015; Ritter-Makinson et al., 2019; Tai et al., 2014). To determine how nRT neuron excitability was affected in this DS mouse model at age P25-P30, we examined intrinsic membrane properties, depolarization-induced spike firing, and hyperpolarization-induced rebound bursting in acute brain slices. The resting membrane potential (RMP) of nRT neurons was significantly reduced in DS mice compared to WT, while input resistance (R_in_), cell capacitance (C_m_), and the membrane time constant (τ) were not significantly altered (**Table 1, Supplementary Figure 1A**). In response to depolarizing current injections, all WT nRT neurons and the majority of DS neurons fired action potentials at a tonic rate, as expected, but 4 of 12 DS neurons fired low-threshold spikes with bursts of action potentials (**Figure 2A**). DS cells were split into two groups, DS tonic and DS burst, for analysis of depolarizing current injections. DS tonic neurons fired significantly fewer action potentials and required significantly more current to elicit an action potential compared to WT neurons (rheobase: WT: 72 ± 14 pA, DS tonic: 118 ± 20 pA; DS burst: 58 ± 13 pA; **Figure 2B,C**). The latency between current injection and the first spike was significantly shorter for DS tonic and longer for DS burst groups compared to WT (WT: 7.4 ± 1.2 ms, DS tonic: 3.8 ± 0.4 ms; DS burst: 16.3 ± 1.9 ms; **Figure 2D**). These data suggest Na_V_1.1 haploinsufficiency alters nRT excitability through impaired depolarization-induced spike firing and a shift toward burst firing mode at rest, which is consistent with the hyperpolarized RMP observed in DS mice.

**Table 1.** Membrane properties of nRT, VPL, and VPM neurons.

|  | nRT |  |  | VPL |  |  | VPM |  |  |
| --- | --- | --- | --- | --- | --- | --- | --- | --- | --- |
|  | WT | DS | P-value | WT | DS | P-value | WT | DS | P-value |
| RMP (mV) | $-51.3 \pm 1.1$ | $-57.1 \pm 1.6$ | *0.018 | $-62.2 \pm 1.3$ | $-69.3 \pm 2.2$ | *0.018 | $-68.8 \pm 1.7$ | $-67.6 \pm 2.6$ | 0.594 |
| C<sub>m</sub> (pF) | $99.8 \pm 0.9$ | $109.5 \pm 6.3$ | 0.270 | $223 \pm 30$ | $207 \pm 28$ | 0.690 | $186 \pm 14$ | $196 \pm 31$ | 0.700 |
| R<sub>in</sub> (MΩ) | $274 \pm 35$ | $254 \pm 33$ | 0.690 | $123 \pm 14$ | $125 \pm 15$ | 0.910 | $116 \pm 10$ | $154 \pm 12$ | *0.030 |
| τ (ms) | $27.4 \pm 3.6$ | $27.0 \pm 3.4$ | 0.930 | $29.8 \pm 6.9$ | $25.9 \pm 4.5$ | 0.660 | $22.0 \pm 3.1$ | $29.8 \pm 5.1$ | 0.190 |
| N | 9 | 15 |  | 12 | 11 |  | 14 | 9 |  |

Intrinsic membrane properties were measured from voltage responses to brief hyperpolarizing current injections. WT and DS values for each parameter were compared by unpaired t-tests. All values are mean ± s.e.m.

**Figure 2.**
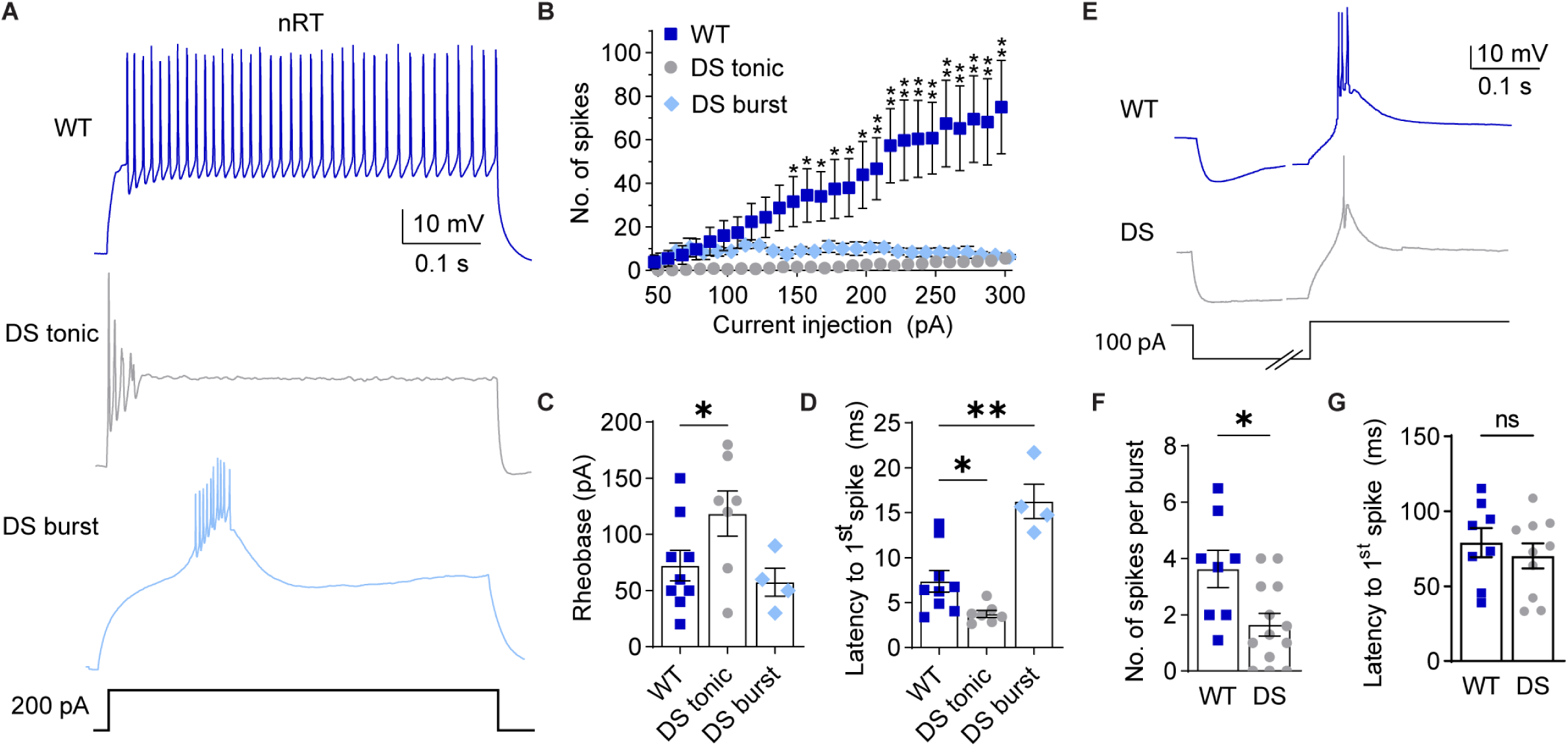
Intrinsic excitability of nRT neurons is altered in DS mice. **A**. Representative traces show nRT neuron spike firing in response to depolarizing current injections. **B**. The number of spikes at each current injection for WT (n = 9), DS neurons that tonically fired (n = 8), and DS neurons that fired bursts (n = 4) were analyzed by mixed-effects analysis for repeated measures (Genotype: p = 0.008, Interaction: p < 0.001) with posthoc Dunnett’s tests at each current injection. *p<0.01 for WT vs. DS tonic. **p< 0.01 for both WT vs DS tonic and WT vs DS burst. **C**. Rheobase and (**D**) latency to the first spike for WT (n = 9), DS tonic (n = 7), and DS burst (n = 4) were analyzed by one-way ANOVA. Rheobase: F(2,17) = 3.812, p = 0.048; posthoc Dunnett’s test, *p = 0.048. Latency: F(2,17) = 20.38, p = 0.001; posthoc Dunnett’s tests, *p = 0.034, **p = 0.015. **E**. Representative traces show rebound burst firing upon recovery from a 500 ms hyperpolarizing current injection. The time axis was broken to facilitate displaying both the hyperpolarization and spike periods. **F**. Unpaired t-tests were used to compare the number of spikes per burst (*p = 0.014, WT: n = 8, DS: n = 13) and (**G**) latency to the first spike (p = 0.489, WT: n = 8, DS: n = 10). The symbols in all bar graphs represent individual neurons.

Hyperpolarization-induced rebound burst firing in nRT neurons is essential for oscillatory activity between nRT and VPL/VPM neurons, and aberrant thalamic oscillations underlie sleep impairments and absence seizures in DS. Therefore, we investigated how Na_V_1.1 haploinsufficiency impacted rebound burst firing after removal of hyperpolarizing current injections in nRT neurons (**Figure 2E**). In DS mice, nRT neurons fired significantly fewer action potentials per burst than WT neurons (WT: 3.6 ± 0.7, DS: 1.6 ± 0.4; **Figure 2F**), with no significant change in burst latency (WT: 79 ± 9.7; DS: 70 ± 8.3 ms; **Figure 2G**). Altogether, these data suggest that Na_V_1.1 haploinsufficiency impairs action potential generation during both tonic and rebound burst firing in nRT neurons.

### Aberrant intrinsic excitability of glutamatergic VPL and VPM neurons in DS mice

Previous studies of the somatosensory thalamus in DS mouse models examined the VB complex, which includes both VPL and VPM neurons; however, these two cell populations have different synaptic connectivity and may have distinct intrinsic membrane properties (Mai and Majtanik, 2018). Therefore, we evaluated how Na_V_1.1 haploinsufficiency impacted the firing properties of these two cell populations independently.

The RMP of VPL neurons was significantly hyperpolarized in DS mice relative to WT, while input resistance, cell capacitance, and membrane time constant were not significantly different (**Table 1, Supplementary Figure 1B**). Consistent with a more hyperpolarized RMP, VPL neurons in DS mice fired significantly fewer spikes than WT mice over a range of depolarizing current injections as detected by a significant interaction effect between genotype and current amplitude in the ANOVA; however, no significant differences were detected by pairwise comparisons at individual current injections (**Figure 3A,B**). In addition, neither the rheobase (WT: 142.9 ± 25.6; DS: 172.2 ± 50.0) nor spike latency (WT: 9.2 ± 1.8 ms; DS: 15.8 ± 3.5 ms) were significantly affected (**Figure 3C,D**). Interestingly, VPL neurons from DS mice fired more spikes per rebound burst upon recovery from hyperpolarization compared to WT (WT: 1.7 ± 0.4 spikes; DS: 3.6 ± 0.6 spikes; **Figure 3F**), while burst latency was not significantly affected (WT: 35.4 ± 5.1 ms; DS: 40.1 ± 3.7 ms; **Figure 3G**). Taken together, these data demonstrate reduced tonic firing and enhanced rebound burst firing in VPL neurons in DS mice, both of which are consistent with the observed hyperpolarized RMP.

**Figure 3.**
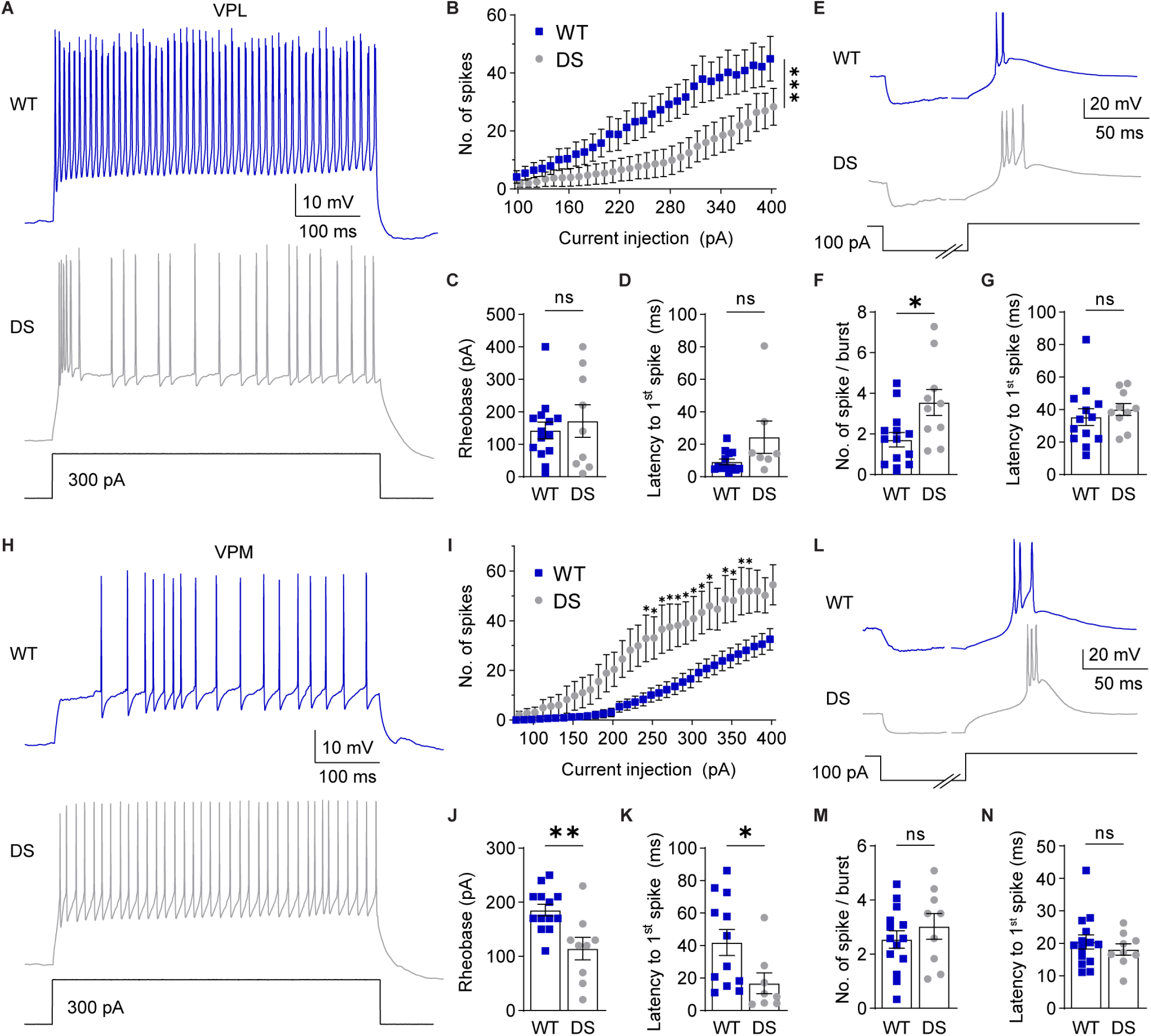
NaV1.1 haploinsufficiency alters intrinsic excitability of VPL and VPM neurons. **A**. Representative traces show VPL spike firing in response to depolarizing current injections. **B**. The number of spikes fired by VPL neurons at each current injection were analyzed by two-way repeated measures ANOVA (WT: n = 13; DS: n = 8; Genotype: p = 0.07, Interaction: ***p < 0.001) with posthoc Sidak’s tests at each current injection (p > 0.05 at each current amplitude). **C**. Rheobase was analyzed by unpaired t-test for WT (n = 14) and DS (n = 9) VPL neurons (p = 0.571). **D**. Latency was analyzed by Mann-Whitney test (p = 0.128) due to failed normality Shapiro-Wilk test. **E**. VPL neuron rebound burst firing upon recovery from hyperpolarization is shown with the time axis broken to facilitate displaying the hyperpolarization and spike periods. **F**. Spikes per burst (*p = 0.01, WT: n = 14, DS: n = 10) and (**G**) burst latency (p = 0.49, WT: n = 13, DS: n = 10) for WT and DS neurons were compared by unpaired t-tests. **H**. Representative traces show VPM spike firing in response to depolarizing current injections. **I**. The number of spikes at each current injection in VPM neurons were analyzed by two-way repeated measures ANOVA (WT: n = 12; DS: n = 9; Genotype: p = 0.011, Interaction: p < 0.001) with posthoc Sidak’s tests at each current injection (*p < 0.01). Unpaired t-tests were used to compare (**J**) rheobase (WT: n = 13; DS: n = 9; **p = 0.004) and (**K**) latency (WT: n = 12; DS: n = 8; *p = 0.037). **L**. Representative traces show rebound burst firing upon recovery from hyperpolarization. Unpaired t-tests were used to compare (**M**) spikes per burst (p = 0.39) and (**N**) burst latency (p = 0.46) for VPM neurons (WT: n = 14, DS: n = 9). The symbols in all bar graphs represent individual neurons.

VPM neurons had significantly greater input resistance in DS mice compared to WT mice, while RMP, cell capacitance, and the membrane time constant were not significantly different (**Table 1, Supplementary Figure 1C**). Given the increased input resistance, VPM neurons would be predicted to have enhanced responses to current injections in DS mice. Indeed, VPM neurons fired significantly more action potentials in DS mice than WT at corresponding depolarizing current injections (**Figure 3H,I**). In addition, DS neurons required significantly less current to elicit an action potential (WT: 185.4 ± 10.8 pA; DS: 114.4 ± 20.8 pA; **Figure 3J**) and had a shorter spike latency (WT: 42.0 ± 8.0 ms; DS: 16.8 ± 6.5 ms; **Figure 3K**). In VPM neurons, the number of spikes per rebound burst (WT: 2.5 ± 0.3; DS: 3.0 ± 0.5) and burst latency (WT: 20.5 ± 2.2 ms; DS: 18.1 ± 1.7 ms) were not significantly different between WT and DS mice (**Figure 3L-N**). Together, these data suggest that VPM neurons have enhanced tonic firing in response to depolarization in DS mice, which may be due to altered intrinsic membrane properties as indicated by increased input resistance.

### Synapse-specific reduction in glutamatergic input to nRT neurons

We hypothesized that altered intrinsic excitability in thalamic neurons would disrupt activity-dependent synapse development and thus exacerbate thalamocortical network dysfunction in DS. The nRT receives glutamatergic inputs from corticothalamic (CT) and thalamocortical (TC) neurons. Deficits in CT-nRT and TC-nRT connectivity would disrupt feed-forward and feed-back inhibition of VPL/VPM neurons, respectively (see **Figure 1A**). To study synapse function in an input-specific manner, we recorded mEPSCs in nRT neurons and took advantage of the distinct decay kinetics of CT-nRT and TC-nRT synaptic currents to separate mEPSCs into two populations (Deleuze and Huguenard, 2016). Faster decaying events, referred to as Type 1, and slower decaying events, referred to as Type 2, have been postulated to represent TC-nRT and CT-nRT synapses, respectively (**Figure 4A,B**).

**Figure 4.**
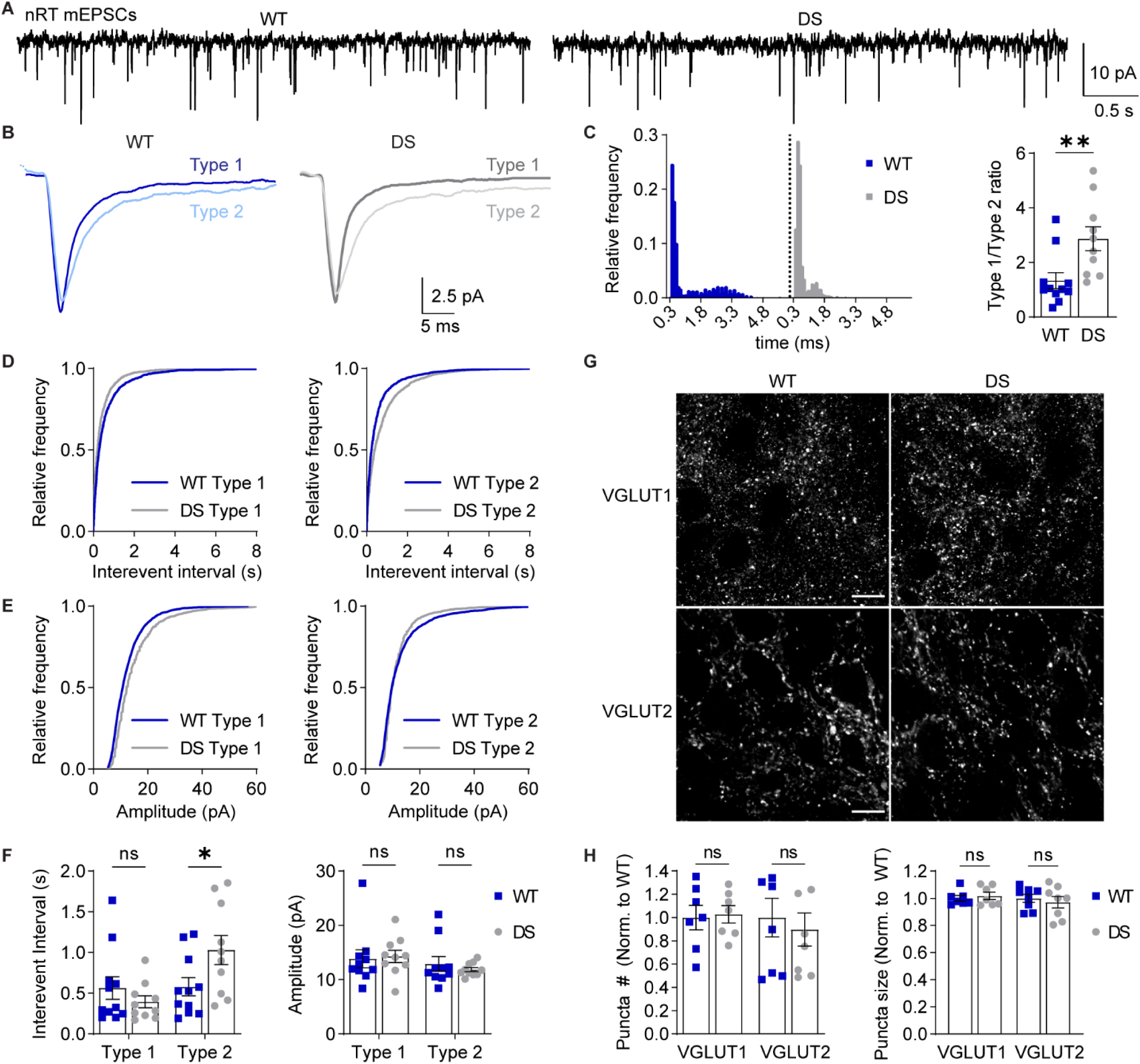
Synapse-specific reduction in nRT mEPSCs in DS mice. **A**. mEPSCs were recorded from nRT neurons in acute brain slices in the presence of TTX. **B**. Representative traces show ensemble averages of WT and DS Type 1 and Type 2 mEPSCs as determined by decay time. **C**. A frequency histogram shows decay times for all events from representative WT and DS neurons. The ratio of Type 1/Type 2 events for WT (n = 11) and DS (n = 10) neurons were analyzed by unpaired t-test (*p = 0.008). Cumulative frequency distributions are shown for **(D)** inter-event interval and **(E)** amplitude of Type 1 and Type 2 mEPSCs from all WT (n = 11) and DS (n = 10) neurons. **F**. Inter-event interval and amplitude values for Type 1 and Type 2 mEPSCs were analyzed by two-way ANOVA with posthoc Sidak’s tests. Inter-event interval: Genotype, F(1,38) = 1.169, p = 0.286; Interaction, F(1,38) = 5.619, p = 0.023; Type 1 WT vs. DS: p = 0.600; Type 2 WT vs DS: *p = 0.039. Amplitude: Genotype, F(1,38) = 0.388, p = 0.538; Interaction, F(1,38) = 0.0699, p = 0.795. **G**. Representative 100X images depict VGLUT1 and VGLUT2 staining in WT and DS nRT. Scale bar: 10 μm. **H**. The number and size of VGLUT1 and VGLUT2 puncta (n = 7) were analyzed by two-way ANOVA. Puncta number: Genotype, F(1,26) = 0.081, p = 0.78; Interaction, F(1,26) = 0.250, p = 0.62. Puncta size: Genotype, F(1,26) = 0.031, p = 0.86; Interaction, F(1,26) = 0.545, p = 0.47. Data points in the bar graphs represent individual neurons (C, F) or mice (H).

Interestingly, the ratio of Type 1 to Type 2 mEPSCs was increased in DS mice relative to WT (WT: 1.3 ± 0.3; DS: 2.9 ± 0.4; **Figure 4C**), with no change in the mean 70-30% decay time for Type 1 events (WT: 2.6 ± 0.3 ms, DS: 3.0 ± 0.4) or Type 2 events (WT: 5.4 ± 0.7 ms; DS: 6.6 ± 0.9 ms; **Supplementary Figure 2**). The inter-event interval was significantly increased for Type 2 events (WT: 0.58 ± 0.11 s; DS: 1.03 ± 0.18 s), while the Type 1 inter-event interval was not significantly altered (WT: 0.56 ± 0.14 s; DS: 0.40 ± 0.08 s; **Figure 4D,F**). There was no main effect of genotype on inter-event interval suggesting that the overall frequency of nRT glutamatergic input was unchanged. Furthermore, mEPSC amplitude was unaltered in both Type 1 (WT: 13.9 ± 1.7 pA, DS: 14.9 ± 1.2 pA) and Type 2 (WT: 12.9 ± 1.3 pA, DS: 11.8 ± 0.4 pA) mEPSC populations in DS mice compared to WT (**Figure 4E,F**). These data suggest that DS mice have a selective decrease in the frequency of Type 2 mEPSCs, which are putative synaptic events at cortical inputs to nRT neurons.

To further evaluate synapse-specific changes in nRT glutamatergic input, we immunostained brain sections from WT and DS mice for CT-specific presynaptic marker VGLUT1 and TC-specific presynaptic marker VGLUT2 (**Figure 4G**). DS mice exhibited no significant differences in either VGLUT1 puncta number (102.8 ± 7.4 % of WT) and size (101.8 ± 2.7 % of WT) or VGLUT2 puncta number (89.7 ± 14.3 % of WT) and size (97.1 ± 4.2 % of WT; **Figure 4H**), suggesting that there is no change in the number of glutamatergic synapses in the nRT. Taken together, the imaging and physiology data are consistent with impaired CT-nRT synapse function, which may result in TC inputs contributing a greater proportion of excitatory synaptic transmission to nRT neurons in this DS mouse model.

In mice aged P25-P30 used herein, there are no GABAergic mIPSCs in nRT neurons and, therefore, we could not evaluate the frequency or strength of spontaneous GABAergic synaptic transmission. However, we did evaluate GABAergic synapse number and size by immunostaining for presynaptic marker VGAT and postsynaptic marker gephyrin in WT and DS tissue. We did not detect any significant differences in VGAT puncta number (102 ± 12% of WT) and size (102 ± 2% of WT), gephyrin puncta number (101 ± 18% of WT) and size (104 ± 2% of WT), or the number (100 ± 10% of WT) and size (103 ± 2% of WT) of VGAT/gephyrin colocalized regions (**Supplementary Figure 3**). If GABAergic synapse density is indeed unchanged in the nRT, then this finding together with the observed reduction in mEPSC frequency might indicate imbalanced excitatory and inhibitory input to the nRT in DS mice.

### Synapse-specific alterations in glutamatergic input to VPL neurons

VPL and VPM neurons receive descending glutamatergic CT inputs as well as ascending glutamatergic sensory inputs through the medial lemniscus and trigeminothalamic pathways, respectively (Lenz, 1992). Previous studies have shown that evoked EPSCs at these two inputs have distinct kinetics, with CT inputs having faster decay times than ascending sensory inputs (Castro-Alamancos, 2002; McCormick and von Krosigk, 1992; Miyata and Imoto, 2006). Therefore, we hypothesized that VPL and VPM mEPSCs could be distinguished based on decay time similarly to nRT neurons. Indeed, a frequency histogram of mEPSC decay times resulted in two distinct populations in VPL (**Figure 5A-C**) and VPM (**Figure 5H-J**) neurons. In both cases, we refer to faster decaying mEPSCs as Type 1 and slower decaying mEPSCs as Type 2.

**Figure 5.**
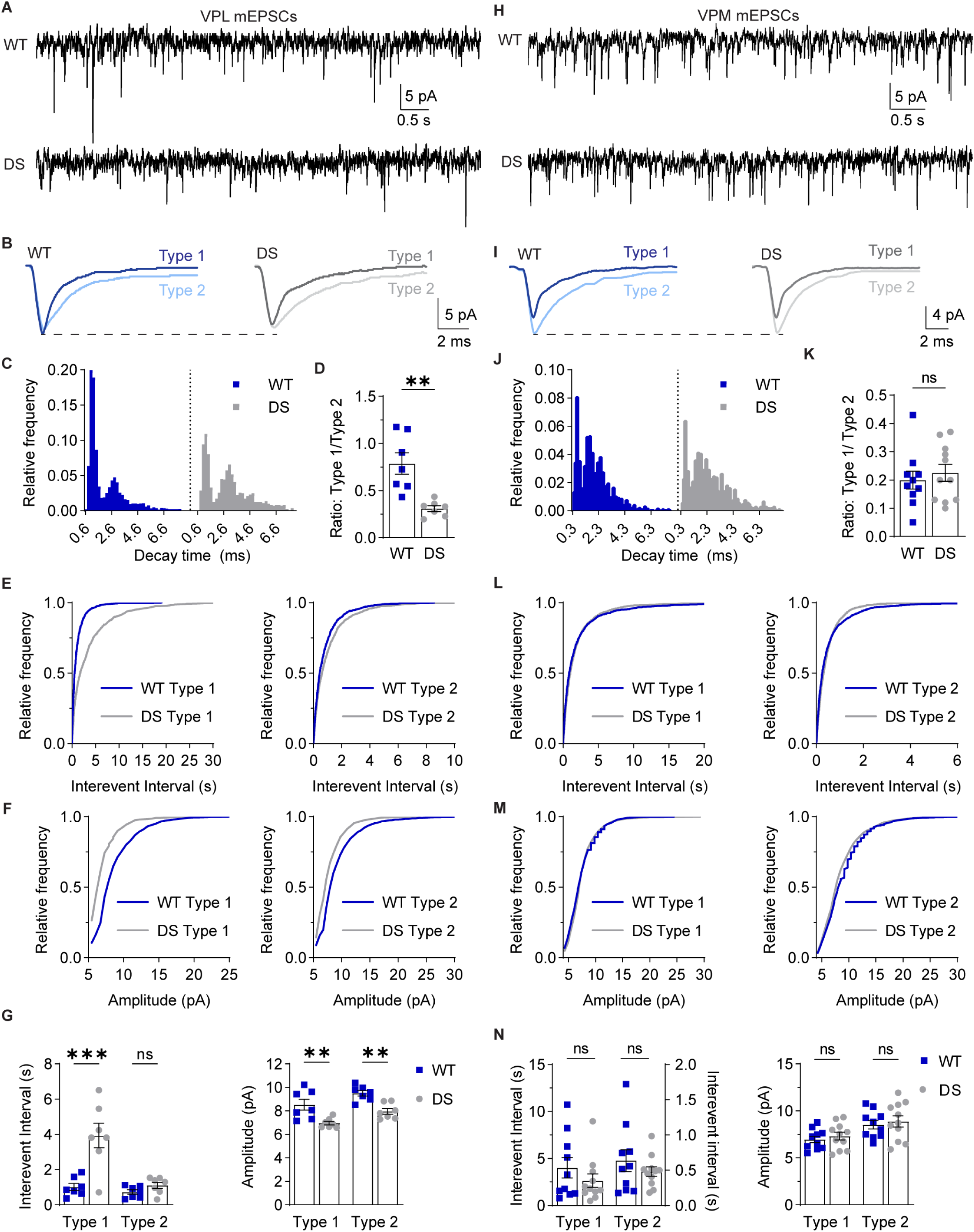
VPL neurons in DS mice exhibit reduced glutamatergic synaptic transmission. **A**. mEPSCs were recorded from VPL and (**H**) VPM neurons in acute brain slices in the presence of 1 μM TTX. **B**. Representative traces show ensemble averages of WT and DS Type 1 and Type 2 mEPSCs from VPL and (**I**) VPM neurons as determined by decay time. **C**. Frequency histogram shows decay times for all events from representative WT and DS VPL neurons. **D**. The ratio of Type 1 to Type 2 events were quantified for each WT and DS neuron (n = 7) and analyzed by unpaired t-test (**p = 0.002). Cumulative frequency distributions are shown for **(E)** inter-event interval and **(F)** amplitude of Type 1 and Type 2 mEPSCs from all WT and DS VPL neurons. **G**. Inter-event interval and amplitude values for Type 1 and Type 2 mEPSCs in VPL neurons were analyzed by two-way ANOVA with posthoc Sidak’s tests. Inter-event interval: Genotype, F(1,24) = 19.28, p < 0.001; Interaction, F(1,24) = 11.42, p = 0.002; Type 1 WT vs. DS: ***p < 0.001; Type 2 WT vs DS: p = 0.73. Amplitude: Genotype, F(1, 24) = 27.15, p < 0.001; Interaction, F(1,24) = 0.002, p = 0.969. Type 1 WT vs. DS: **p = 0.002; Type 2 WT vs DS: **p = 0.003. **J**. The frequency histogram shows decay times for all events from representative WT and DS VPM neurons. **K**. Type 1/Type 2 event ratios for WT (n = 10) and DS (n = 11) VPM neurons were analyzed by unpaired t-test (p = 0.57). Cumulative frequency distributions are shown for **(L)** inter-event interval and **(M)** amplitude of Type 1 and Type 2 mEPSCs from all WT and DS VPM neurons. **N**. Inter-event interval and amplitude values for Type 1 and Type 2 mEPSCs in VPM neurons were analyzed by two-way ANOVA with posthoc Sidak’s tests. Inter-event interval: Genotype, F(1,38) = 1.407, p = 0.243; Interaction, F(1,38) = 0.893, p = 0.351; Type 1 WT vs. DS: p = 0.261; Type 2 WT vs DS: p = 0.982. Amplitude: Genotype, F(1, 38) = 0.573, p = 0.454; Interaction, F(1,38) < 0.001, p = 0.993. Type 1 WT vs. DS: p = 0.840; Type 2 WT vs DS: p = 0.833. Data points in the bar graphs represent individual neurons.

In VPL neurons, the ratio of Type 1 to Type 2 mEPSCs was decreased in DS mice relative to WT (WT: 0.79 ± 0.11; DS: 0.31 ± 0.03; **Figure 5D**), with no change in the decay times for Type 1 mEPSCs (WT: 5.9 ± 0.8 ms; DS: 6.2 ± 0.8 ms) or Type 2 mEPSCs (WT: 9.2 ± 0.8 ms; DS: 8.8 ± 1.0 ms; **Supplementary Figure 4A,B**). The change in relative proportions of Type 1 and Type 2 mEPSCs was driven by a significant increase in the inter-event interval for Type 1 mEPSCs (WT: 1.01 ± 0.20 s; DS: 3.95 ± 0.69 s), whereas the Type 2 mEPSC inter-event interval was not significantly affected (WT: 0.73 ± 0.12 s; DS: 1.11 ± 0.17 s; **Figure 5E,G**). Furthermore, mEPSC amplitude was significantly reduced for both Type 1 (WT: 8.5 ± 0.5 pA; DS: 7.0 ± 0.2 pA) and Type 2 (WT: 9.5 ± 0.2 pA; DS: 8.0 ± 0.2 pA) events in DS VPL neurons compared to WT (**Figure 5F,G**).These data suggest that VPL neurons in DS mice have a global decrease in glutamatergic synapse strength and an input-specific reduction in ascending glutamatergic synaptic inputs.

Interestingly, the ratio of Type 1 and Type 2 events in VPM neurons was not significantly different between WT and DS mice (WT: 0.2 ± 0.03; DS: 0.23 ± 0.03; **Figure 5K**), and mEPSC decay times were similar for Type 1 (WT: 3.5 ± 0.3 ms; DS: 4.0 ± 0.5 ms) and Type 2 mEPSCs (WT: 7.0 ± 0.7 ms; DS: 8.0 ± 0.6 ms; **Supplementary Figure 4C,D**). VPM neurons in WT and DS mice also exhibited no significant differences in the inter-event interval of Type 1 (WT: 4.0 ± 1.1 s; DS: 2.6 ± 0.7 s) and Type 2 mEPSCs (WT: 0.64 ± 0.16 s; DS: 0.48 ± 0.07 s) or the amplitude of Type 1 (WT: 6.9 ± 0.3 pA; DS: 7.3 ± 0.4 pA) and Type 2 mEPSCs (WT: 8.5 ± 0.5 pA; DS: 8.9 ± 0.6 pA; **Figure 5L-N**). Taken together, the mESPC data suggests that Na_V_1.1 haploinsufficiency may effect synapse development in the VPL and VPM neurons differently leading to cell-type-specific roles for these glutamatergic neuron populations in thalamic dysfunction in DS.

To further evaluate glutamatergic synapses in the VPL and VPM, we immunostained for VGLUT1 to label CT synapses and VGLUT2 to label ascending sensory synapses, and then quantified the number and size of synaptic puncta (**Figure 6A,B**). The number of VGLUT2 puncta in the VPL was decreased in DS mice relative to WT (83.7 ± 3.2% of WT), whereas the number of VGLUT1 puncta was unchanged (99.9 ± 6.7% of WT, **Figure 6C**). The size of VGLUT1 and VGLUT2 puncta were not significantly different between WT and DS mice (VGLUT1: 96.7 ± 3.2 % of WT; VGLUT2: 107 ± 6.1 % of WT). Furthermore, the VPM in WT and DS mice had similar VGLUT1 and VGLUT2 puncta number (VGLUT1: 1.12 ± 5.9 % of WT; VGLUT2: 87.0 ± 12.7 % of WT) and size (VGLUT1: 96.2 ± 3.2 % of WT; VGLUT2: 101.5 ± 7.2 % of WT; **Figure 6D**). Altogether, the imaging and electrophysiological findings are consistent with a synapse-specific reduction in the number of ascending sensory synapses in the VPL as well as a reduction in overall glutamatergic synapse strength specifically in VPL neurons.

**Figure 6.**
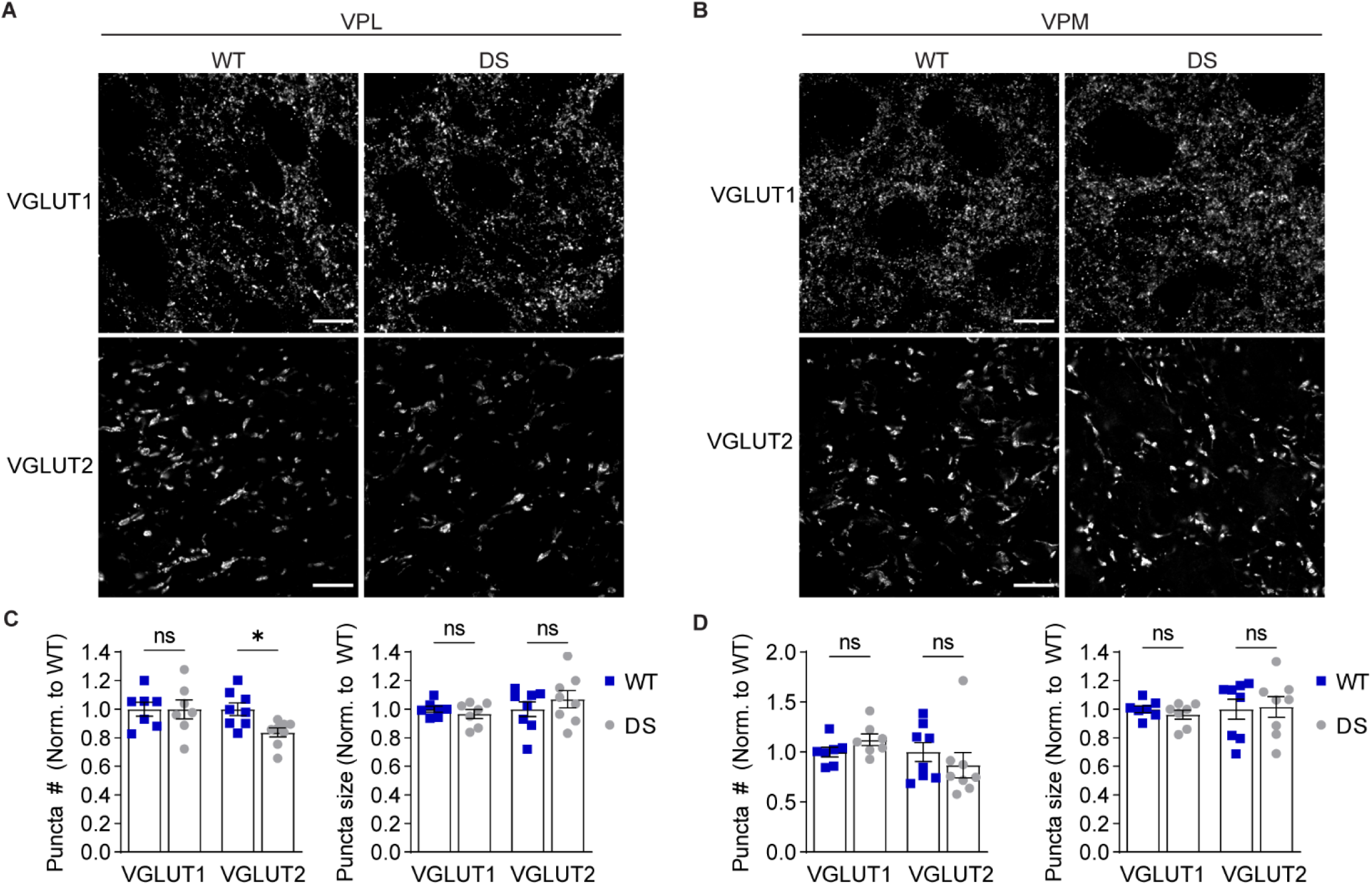
DS mice exhibit reduced ascending sensory input to the VPL. Representative 100X images show VGLUT1 (scale bar: 10 μm) and VGLUT2 (scale bar: 20 μm) immunostaining in the (**A**) VPL and (**B**) VPM. The number and size of VGLUT1 and VGLUT2 puncta were quantified for the (**C**) VPL and (**D**) VPM (n = 7) and analyzed by two-way ANOVA. VPL puncta number: Genotype, F(1,26) = 2.914, p = 0.10; Interaction, F(1,26) = 2.829, p = 0.10; posthoc Sidak’s tests: VGLUT1, p = 0.99; VGLUT2, *p = 0.04. VPL puncta size: Genotype, F(1,26) = 0.161, p = 0.69; Interaction, F(1,26) = 1.262, p = 0.27; posthoc Sidak’s tests: VGLUT1, p = 0.86; VGLUT2, p = 0.47. VPM puncta number: Genotype, F(1,26) = 0.002, p = 0.96; Interaction, F(1,26) = 1.9, p = 0.18; posthoc Sidak’s tests: VGLUT1, p = 0.60; VGLUT2, p = 0.52. VPM puncta size: Genotype, F(1,26) = 0.040 p = 0.84; Interaction, F(1,26) = 0.221, p = 0.64; posthoc Sidak’s tests: VGLUT1, p = 0.88; VGLUT2, p = 0.98. Data points in all bar graphs represent individual mice.

### Inhibitory synaptic input is enhanced in VPL neurons

VPL and VPM neurons receive inhibitory input from nRT neurons, with VPL neurons receiving input from both somatostatin- and parvalbumin-positive neurons and VPM neurons receiving input from parvalbumin-positive nRT neurons (Clemente-Perez et al., 2017). We hypothesized that aberrant excitability within the somatosensory thalamic circuitry would alter inhibitory synaptic input from the nRT to VPL and VPM neurons. Therefore, we analyzed the decay time, frequency, and amplitude of mIPSCs in VPL and VPM neurons (**Figure 7A,G**). VPL neuron mIPSC decay time was significantly faster in DS mice (16.8 ± 2.4 ms) compared to WT (25.4 ± 2.1 ms; **Figure 7B**), which suggests that either postsynaptic receptor composition or the proximal-distal distribution of GABAergic inputs is altered in DS mice. Moreover, VPL neurons exhibited reduced mIPSC inter-event interval in DS mice (0.132 ± 0.023 ms) compared to WT (0.266 ± 0.033 ms), whereas VPL mIPSC amplitude was not significantly altered in DS mice (23.0 ± 1.1 pA) relative to WT (20.3 ± 0.6 pA; **Figure 7C,D**). VPM neurons in DS mice had no significant differences relative to WT in mIPSC decay time (WT: 29.8 ± 5.5 ms; DS: 35.8 ± 7.9 ms), inter-event interval (WT: 0.294 ± 0.06 s; DS: 0.269 ± 0.06 s), or amplitude (WT: 18.2 ± 0.7 pA; DS: 19.7 ± 0.6 pA; **Figure 7H-J**). These data suggest that Na_V_1.1 haploinsufficiency enhances the frequency of GABAergic input to VPL neurons without affecting GABAergic input to VPM neurons.

**Figure 7.**
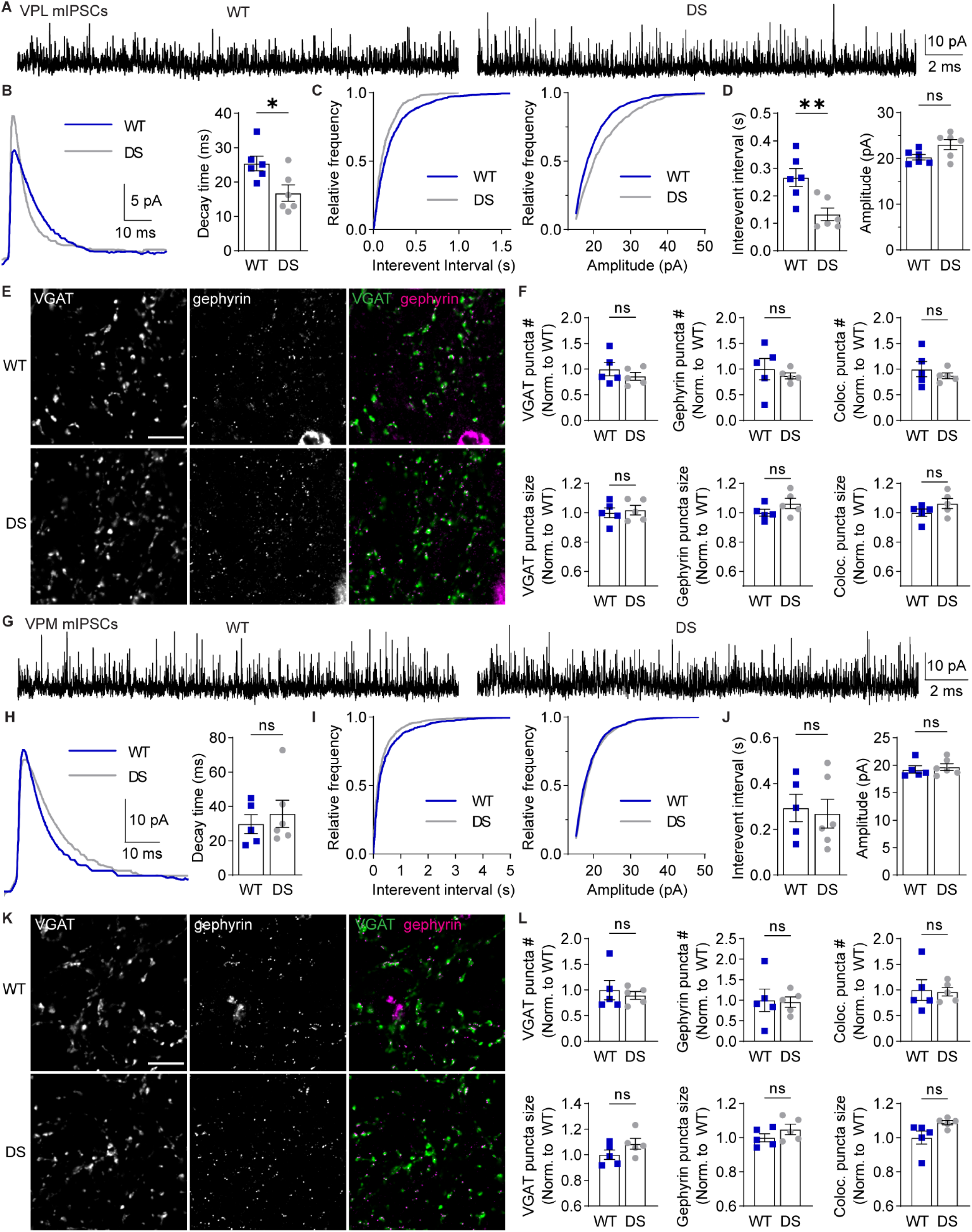
VPL neurons in DS mice exhibit reduced GABAergic synaptic transmission. **A**. mIPSCs were recorded from VPL neurons in acute brain slices in the presence of 1 μM TTX. **B**. Representative traces show ensemble averages of WT and DS mIPSCs, which were were fitted to determine decay times for WT and DS neurons (n = 6) and analyzed by unpaired t-test (*p = 0.02). **C**. Cumulative frequency distributions are shown for mIPSC inter-event interval and amplitude for all WT and DS neurons. **D**. Inter-event interval and amplitude for each WT and DS neuron were quantified and analzyed by unpaired t-tests. Inter-event interval: **p = 0.008. Amplitude: p = 0.054. **E**. Representative 100X images show VGAT (scale bar: 10 μm) and gephyrin immunolabeling in the VPL. **F**. The number and size of VGAT, gephyrin, and VGAT-gephryin colocalized puncta in the VPL (n = 5) were compared by unpaired t-tests. VGAT puncta number: p = 0.39, size: p = 0.72; gephyrin puncta number: p = 0.39, size p = 0.72; colocalized puncta number: p = 0.44, size: p = 0.18. **G**. mIPSCs were recorded from VPM neurons, and (**H**) representative traces show ensemble averages of WT and DS mIPSCs, which were fitted to determine decay time for WT (n = 5) and DS (n = 6) neurons and analyzed by an unpaired t-test (p = 0.57). **I**. Cumulative frequency distributions were plotted for mIPSC inter-event interval and amplitude. **J**. Group data were analyzed by unpaired t-tests. Inter-event interval: p = 0.785. Amplitude: p = 0.635. **K**. Representative 100X images show VGAT (scale bar: 10 μm) and gephyrin immunolabeling in the VPM. **L**. The number and size of VGAT, gephyrin, and VGAT-gephryin colocalized puncta were quantified in the VPM (n = 5) and compared by unpaired t-tests. VGAT puncta number: p = 0.62, size: p = 0.16; gephyrin puncta number: p = 0.89, size p = 0.24; colocalized puncta number: p = 0.88, size: p = 0.06. The data points in the bar graphs represent individual neurons (B, D, H, J) or mice (F, L).

To further evaluate how GABAergic inputs were altered in DS mice, we immunolabeled VGAT and gephyrin in WT and DS thalamus and quantified the number and size of inhibitory synapses. We detected no significant differences in WT and DS VPL thalamus when comparing VGAT puncta number (86 ± 7.5 % of WT), VGAT puncta size (102 ± 3.2 % of WT), gephyrin puncta number (86 ± 7.5 % of WT), or gephyrin puncta size (102 ± 3.2 % of WT; **Figure 7E,F**). We also quantified colocalized regions of VGAT and gephyrin puncta in the VPL and found no significant differences in the number (87 ± 5.6 % of WT) or size (106 ± 3.4 % of WT; **Figure 7E**). Similarly, we did not find any significant differences in GABAergic synapse labeling in VPM when we compared VGAT puncta number (90 ± 7.5 % of WT), VGAT puncta size (109 ± 4.2 % of WT), gephyrin puncta number (96 ± 12.4 % of WT), or gephyrin puncta size (105 ± 3.0 % of WT; **Figure 7K,L**). We also found no significant differences in colocalized VGAT and gephyrin puncta number (97 ± 8.6 % of WT) or size (108 ± 1.3 % of WT) in the VPM (**Figure 7L**). Taken together, the mIPSC and VGAT/gephyrin data are consistent with enhanced frequency and accelerated kinetics of GABAergic synaptic transmission in VPL neurons of DS mice, which may occur through changes in presynaptic release and/or postsynaptic receptor expression as opposed to changes in GABAergic synapse number.

## DISCUSSION

Collectively, these data provide novel insight into the mechanisms underlying somatosensory CT circuit dysfunction in a Na_V_1.1 haploinsufficiency DS mouse model. The intrinsic excitability of GABAergic nRT neurons and glutamatergic VPL and VPM neurons was disrupted in a cell-type-specific manner, including alterations in both depolarization-induced tonic firing and hyperpolarization-induced rebound burst firing in nRT and VPL neurons. Unexpectedly, VPL and VPM neurons exhibited opposing changes to depolarization-induced firing, suggesting they may differentially contribute to circuit-wide dysfunction. Glutamatergic synaptic input to both the nRT and VPL was reduced in an input-specific manner, and GABAergic synaptic input to the VPL was enhanced, further contributing to an imbalance of synaptic input to the region. Together, these results indicate that synaptic- and cellular-level changes in GABAergic and glutamatergic neuron populations contribute to somatosensory thalamic dysfunction in DS.

As expected, Na_V_1.1 haploinsufficiency impaired both tonic and burst firing in GABAergic nRT neurons. These changes in nRT excitability are consistent with previous findings of reduced depolarization-induced tonic firing and hyperpolarization-induced rebound bursts in *Scn1a*-haploinsufficient and Na_V_1.1-R1648H mouse models of DS (Hedrich et al., 2014; Kalume et al., 2015). While we observed a reduction in the number of spikes per burst in nRT DS neurons, we additionally found nRT neurons exhibited a hyperpolarized RMP and approximately 30% responded to depolarization with burst firing, suggesting they may have a greater propensity to be in a burst firing mode. In contrast to our findings, the Na_V_1.1-R1407X mouse model exhibited hyperexcitability of parvalbumin-positive nRT neurons including prolonged hyperpolarization-induced rebound bursts and enhanced depolarization-induced spike firing (Ritter-Makinson et al., 2019). Discrepancies between models could be attributed to the specific etiology of the disease model or the background strain as DS models are known to have a high degree of strain-dependent phenotype variability (Miller et al., 2014; Rubinstein et al., 2015b). The age of the mice could also cause discrepancies between studies as some cortical neuron populations exhibit a transient window of altered excitability (Favero et al., 2018); however, developmental studies of thalamic dysregulation have not been reported in DS mouse models. From a therapeutic perspective, these model-specific effects indicate that successful patient treatment may largely depend on the patient’s specific genetic alteration. As truncation mutations resulting in haploinsufficiency are responsible for about half of all DS cases, the mechanisms uncovered in this study may be relevant to a large population of patients with DS.

A novel discovery of this study is the opposing changes in the excitability of glutamatergic thalamocortical neurons in the VPL and VPM as well as distinct effects on intrinsic membrane properties. Many glutamatergic neuron populations express Na_V_1.1 at low levels and reportedly exhibit no altered excitability in DS, but recent evidence suggests that CA1 pyramidal neuron excitability is altered and changes over the time course of the disease (Almog et al., 2021). Interestingly, previous investigations have reported no alterations in the excitability of VB neurons in DS models (Hedrich et al., 2014; Ritter-Makinson et al., 2019). This could be due to differences in the specific DS model, the developmental time period studied, or a lack of differentiating between the VPL and VPM, the two regions comprising the VB thalamus, which could have resulted in an averaging of cell-type-specific alterations and no overall change.

The cellular properties of VPL and VPM neurons have not yet been directly compared, and thus our understanding of how Na_V_1.1 haploinsufficiency leads to opposing changes in their excitability is limited. We speculate that different Na_V_1.1 expression levels or subcellular localization may underlie distinct roles for Na_V_1.1 in normal VPL and VPM neuron physiology. Alternatively, the expression of different complements of Na_V_, Ca_V_, or K_V_ channels in VPL and VPM neurons could lead to distinct compensatory mechanisms that underlie cell-type-specific alterations to membrane properties in the DS model. This is supported by evidence from a spinal cord injury model wherein VPL neurons exhibit increased Na_V_1.3 expression post-injury, while VPM neurons do not (Zhao et al., 2006). Additionally, reduced tonic glutamatergic input to VPL neurons could hyperpolarize their RMP, thereby contributing to hypoexcitability. Regardless of the specific mechanism leading to distinct changes in intrinsic properties, these findings suggest that the VPL and VPM may contribute differentially to circuit dysfunction and successful correction of circuit function may require cell-type-specific therapeutic targets.

An additional discovery of this study is altered synaptic connectivity in the thalamus. Both nRT and VPL neurons in this mouse model exhibited an input-specific reduction in mEPSCs, and VPL neurons exhibited an augmentation of mIPSC frequency. The Na_V_1.1-R1648H DS model previously showed reduced spontaneous IPSCs in GABAergic nRT and glutamatergic thalamocortical neurons, but no changes in mIPSCs (Hedrich et al., 2014). Another DS model exhibited no change in spontaneous EPSCs in nRT or VB neurons (Ritter-Makinson et al., 2019). These discrepancies could be due to age-dependent changes in synaptic connectivity and the specific cell populations that were studied. If impaired intrinsic excitability early in the disease leads to subsequent alterations in synapse formation and maintenance, studies at earlier time points in development may not reveal such synaptic-level changes. Developmental studies would be required to elucidate the time course over which intrinsic and extrinsic alterations occur.

Both the cell-type- and input-specific nature of these synaptic alterations suggests that tuning circuit excitability may require pathway-specific therapeutic targets. The distinct changes in VPL and VPM synaptic connectivity could be due to their respective changes in excitability or their distinct sources of ascending input as the VPL and VPM receive somatosensory information from the body and head, respectively. Indeed, the observed reduction in VGLUT2-positive synapses as well as the frequency of Type 1 mEPSCs (the population with faster decay times) in the VPL supports a specific reduction in ascending sensory inputs (Miyata and Imoto, 2006). A most intriguing finding is the reduced CT input to nRT neurons without a corresponding reduction in CT input to either VPL or VPM neurons. The CT axons that innervate nRT neurons are collaterals of those innervating VPL and VPM neurons, suggesting a postsynaptic mechanism specific to CT-nRT synapses underlies altered synapse development and/or function. We did not detect a corresponding change in VGLUT1 puncta in the nRT, which is consistent with altered glutamate release; however, given that CT neurons do not express Na_V_1.1 highly, the data rather implicate a postsynaptic mechanism such as altered glutamate receptor expression or dendritic filtering. Determining the underlying mechanism would require investigating specific glutamate receptor populations as well as studying synaptic potentials under more physiological conditions in addition to the voltage-clamp studies conducted herein.

From a clinical perspective, the dysfunction in VPL and VPM neurons observed here may present a unique opportunity for therapeutic intervention. Specifically, the diversity of glutamatergic receptor expression in the thalamus provides a wide array of therapeutic targets. Modulating glutamate receptors is becoming increasingly feasible as a variety of subtype-selective modulators for metabotropic glutamate receptors, AMPA receptors, and NMDA receptors are being developed (Azumaya et al., 2017; Hansen et al., 2018; Hovelso et al., 2012; Kadriu et al., 2021; Mazzitelli et al., 2018). For example, a recent study indicated that the GluN2A positive allosteric modulator GNE-0723 reduces low-frequency oscillations and epileptiform activity in a DS mouse model, providing evidence that NMDA receptor modulation could be a viable therapeutic option (Hanson et al., 2020). The GluN2C and GluN2D subunits of NMDA receptors have more restricted expression patterns, including relatively high expression in the thalamus, and recently developed GluN2C/2D-selective modulators could offer a means to tune thalamic function with more limited adverse effects (Acker et al., 2011; Khatri et al., 2014; Liu et al., 2019; Mullasseril et al., 2010; Swanger et al., 2018; Yi et al., 2020). Thus, the data presented here lay the foundation to explore glutamatergic synapse modulation as a possible therapeutic strategy to correct thalamic dysfunction in DS.

This evidence regarding cell-type-specific dysfunction advances our understanding of how GABAergic and glutamatergic neuron populations may together contribute to pathological thalamocortical network function (see model in **Figure 8**). We posit that altered depolarization-induced tonic firing may impair VPL cell output and enhance VPM cell output in response to ascending sensory information. The reduced ascending input to the VPL may further impair somatic information processing, perhaps contributing to reduced sensitivity to pain and attention deficits evident in the disease (**Figure 8A,B**) (Catarino et al., 2011; Villas et al., 2017). Burst firing in the somatosensory thalamus underlies oscillatory activity critical for thalamocortical network function, and disrupted burst firing is associated with absence seizures, sleep disorders, and chronic pain (Fogerson and Huguenard, 2016; Hains et al., 2006; Henderson et al., 2013). Intra-thalamic oscillations occur when nRT neurons at a hyperpolarized RMP receive depolarizing input from the cortex, which leads to feed-forward nRT-VPL and nRT-VPM inhibition. VPL and VPM neurons are then hyperpolarized and will burst upon recovery from hyperpolarization leading to feedback inhibition. In this DS model, VPL neurons had a hyperpolarized RMP and fired significantly more spikes during burst firing. The nRT neurons exhibited a hyperpolarized RMP and an increased propensity to burst in response to depolarization, which could result in augmented feed-forward inhibition of VPL and VPM neurons (**Figure 8A,C**). Thus, together these mechanisms may result in enhanced bursting in all three cell populations that contributes to aberrant thalamic oscillations in DS (Kalume et al., 2015; Ritter-Makinson et al., 2019). A complete understanding of how intrinsic and synaptic mechanisms contribute to circuit dysfunction will require elucidating how cells respond to excitatory drive from ascending and descending synaptic inputs.

**Figure 8.**
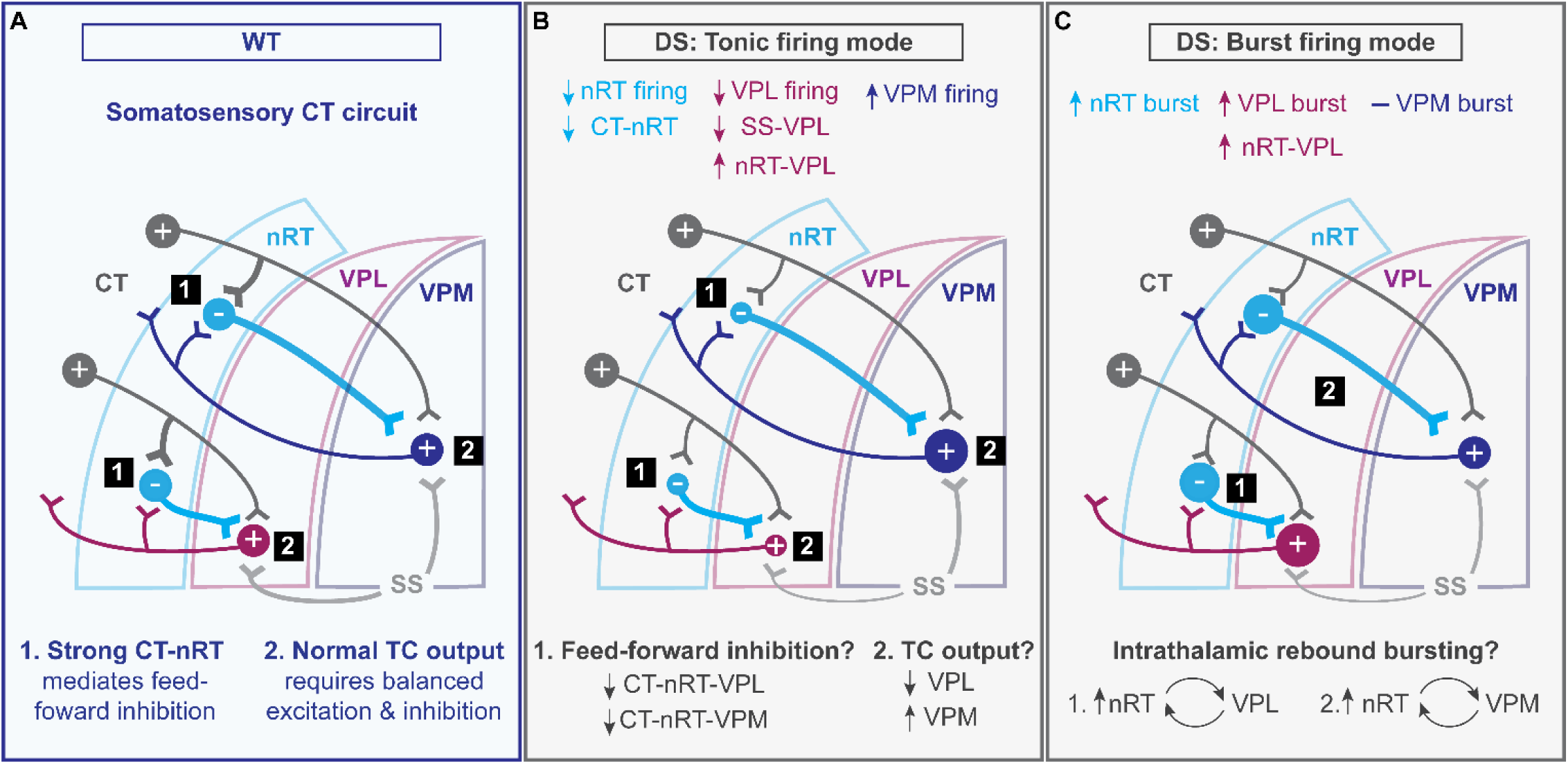
Hypothetical model of somatosensory CT circuit dysfunction in DS. **A**. In WT mice, normal somatosensory CT circuit function involves (1) feed-forward inhibition that requires a strong CT-nRT connection, which drives nRT inhibition of VPL and VPM neurons during both tonic sensory processing and oscillatory burst firing modes, and (2) a critical balance of excitatory and inhibitory synaptic input that controls thalamocortical (TC) output from the VPL and VPM in response to ascending somatosensory (SS) excitation. **B**. In DS mice, we observed that nRT neurons had reduced tonic firing and CT-nRT excitation, VPL neurons had reduced firing, reduced ascending SS excitation, and increased nRT-VPL inhibition, and VPM neurons had enhanced tonic firing. In tonic firing mode, we posit that these effects would combine to reduce feed-forward inhibition of both VPL and VPM, reduce VPL TC output (due to reduced firing and SS input), and enhance VPM TC output. **C**. Regarding burst firing in DS mice, we observed a shift towards burst firing in nRT neurons at rest, although fewer spikes were fired per burst, and enhanced burst firing in VPL neurons, which may enhance nRT –VPL and nRT –VPM intrathalamic rebound burst firing.

## Conclusions

In summary, this work discovered cell-type-specific dysregulation of synapses and intrinsic excitability within the somatosensory thalamus in a DS mouse model. Specifically, the findings introduce altered glutamatergic neuron excitability and synapse function as disease mechanisms that may contribute to thalamocortical network dysfunction underlying aberrant sensory processing, disrupted sleep, and absence seizures. The cell-type-specific intrinsic and synaptic disease mechanisms could affect thalamic function distinctly and thus contribute to particular symptoms or developmental stages of DS. Further investigation is required to elucidate how these previously unidentified mechanisms contribute to DS symptomology across the course of the disease and whether modulating specific therapeutic targets in the glutamate system can restore thalamic function and ameliorate corresponding phenotypes.

## Abbreviations

DS: Dravet syndrome
nRT: reticular nucleus of the thalamus
VPM: ventral posteromedial nucleus
VPL: ventral posterolateral nucleus
VB: ventrobasal complex
CT: corticothalamic
TC: thalamocortical

## Author contributions

*Carleigh Studtmann:* Validation, Formal analysis, Investigation, Writing – original draft, Visualization; *Marek Ladislav:* Methodology, Validation, Writing – review & editing, Supervision; *Mackenzie A. Topolski:* Validation, Investigation, Writing – review & editing; *Mona Safari:* Investigation, Writing – review & editing; *Sharon A. Swanger:* Conceptualization, Methodology, Validation, Formal analysis, Investigation, Resources, Writing – original draft, Visualization, Supervision, Project administration, Funding acquisition.

