## Supplementary Data for "Synaptic and intrinsic mechanisms impair reticular thalamus and thalamocortical neuron function in a Dravet syndrome mouse model"

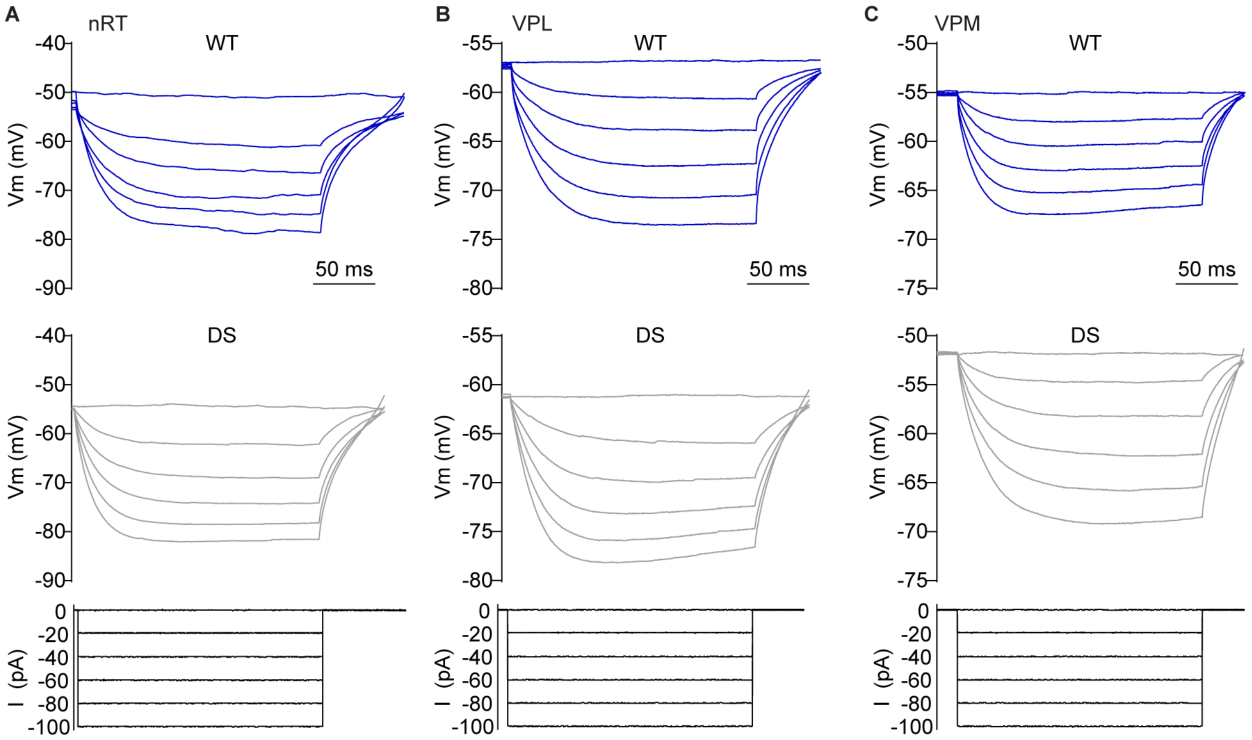


**Supplementary Figure 1. Recordings for intrinsic membrane properties of nRT, VPL, and VPM neurons.** Representative current-clamp recordings show voltage responses to 200 ms hyperpolarizing current injections for (**A**) nRT, (**B**) VPL, and (**C**) VPM neurons from WT and DS mice. The bottom panels show current injection amplitude.

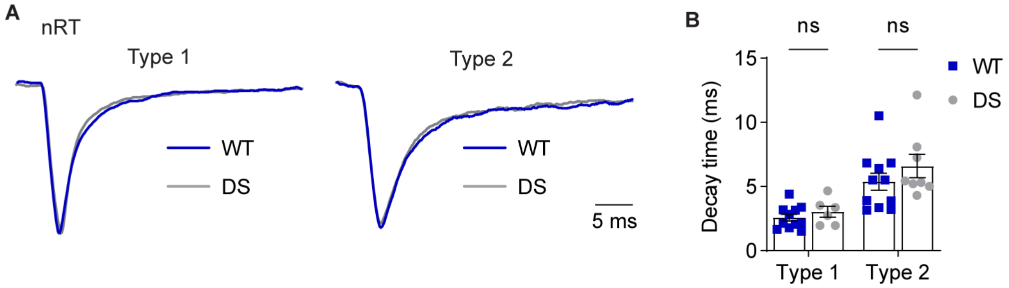


**Supplementary Figure 2. mEPSC decay times do not differ between WT and DS nRT neurons. A.** Traces are ensemble averages of Type 1 and Type 2 mEPSCs from representative WT and DS neurons. Traces were normalized to the respective WT mEPSC amplitude. **B.** The decay times were measured by fitting the ensemble average of Type 1 and Type 2 mEPSCs for each WT and DS neuron. Group data were plotted as mean ± s.e.m. and analyzed by two-way ANOVA. Genotype: $F(1,32) = 1.740$, $p = 0.197$; mEPSC Type: $F(1,32) = 25.02$, $p < 0.001$; Interaction: $F(1,32) = 0.359$, $p = 0.553$.

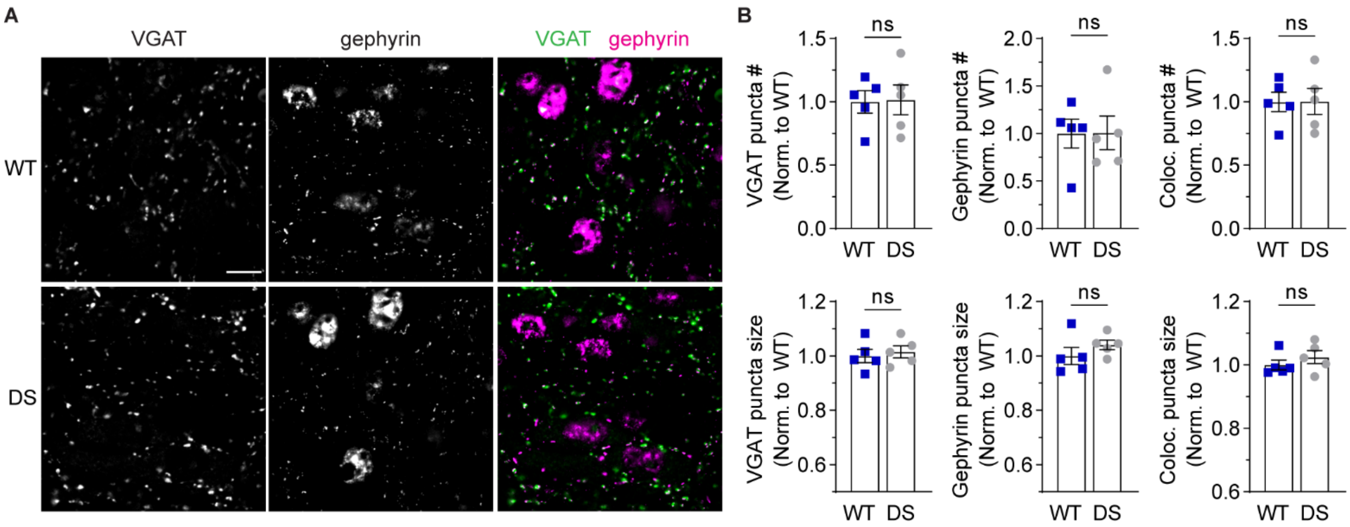


**Supplementary Figure 3. GABAergic synapse number and size in the nRT are unchanged in DS mice. A.** Representative 100X images show VGAT (scale bar: 10 µm) and gephyrin immunolabeling in the nRT as well as the merged image. **B.** The number and size of VGAT, gephyrin, and vGat-gephryin colocalized puncta were quantified, plotted as group mean ± s.e.m. with data points representing individual mice (n = 5), and compared by unpaired t-tests. VGAT puncta number: p = 0.91, size: p = 0.66; gephyrin puncta number: p = 0.98, size p = 0.29; colocalized puncta number: p = 0.98, size: p = 0.36.

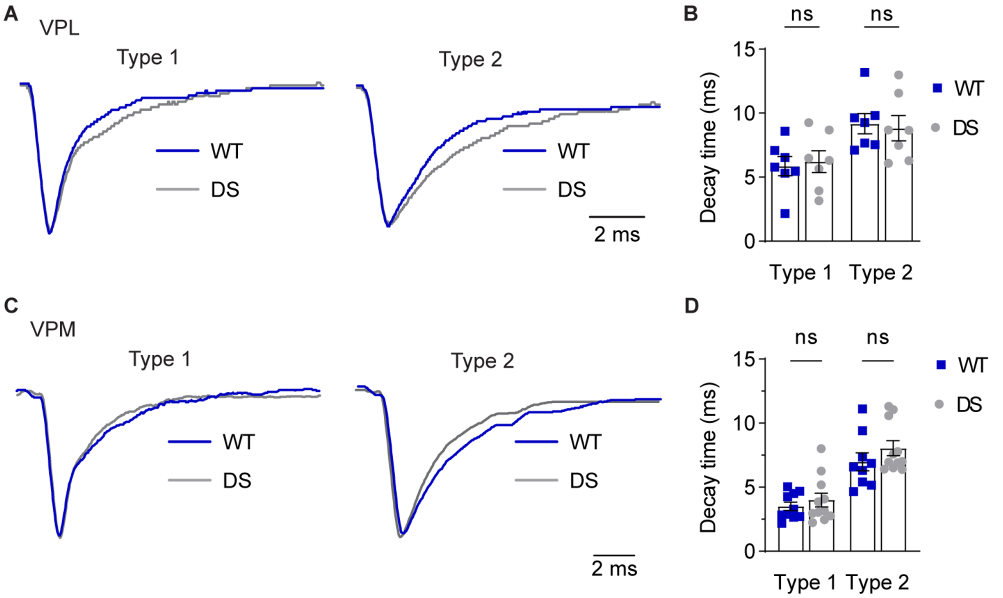


**Supplementary Figure 4. VPL and VPM mEPSC decay times are similar between WT and DS mice.** Traces are ensemble averages of Type 1 and Type 2 mEPSCs from representative (**A**) VPL and (**C**) VPM neurons. Traces were normalized to the respective WT mEPSC amplitude. The decay times were measured by fitting the ensemble average of Type 1 and Type 2 mEPSCs for each WT and DS neuron. **B,D**. Group data were plotted as mean ± s.e.m. and analyzed by two-way ANOVA. VPL: Genotype, $F(1,24) = 0.001$ $p > 0.99$; mEPSC Type, $F(1,24) = 12.22$, $p = 0.002$; Interaction, $F(1,24) = 0.178$, $p = 0.68$. VPM: Genotype, $F(1,37) = 0.261$, $p = 0.612$; mEPSC Type, $F(1,37) = 46.69$, $p < 0.001$; Interaction, $F(1,37) = 1.987$, $p = 0.167$.
